# An exploration of mechanisms underlying *Desemzia incerta* colonization resistance to methicillin-resistant *Staphylococcus aureus* on the skin

**DOI:** 10.1101/2023.10.11.561853

**Authors:** Monica Wei, Simon AB Knight, Hossein Fazelinia, Lynn Spruce, Jennifer Roof, Emily Chu, Jasmine Walsh, Laurice Flowers, Daniel Y. Kim, Jun Zhu, Elizabeth A. Grice

**Affiliations:** University of Pennsylvania, Perelman School of Medicine, Department of Dermatology and Microbiology, Philadelphia, Pennsylvania, USA; Children’s Hospital of Philadelphia, Proteomics Core Facility, Philadelphia, Pennsylvania, USA

## Abstract

Colonization of human skin and nares by methicillin-resistant *Staphylococcus aureus* (MRSA) leads to community spread of MRSA. This spread is exacerbated by transfer of MRSA between humans and livestock, particularly swine. Here we capitalized on the shared features between human and porcine skin, including shared MRSA colonization, to study novel bacterial mediators of MRSA colonization resistance. We focused on the poorly studied bacterial species *Desemzia incerta*, which we found to exert antimicrobial activity through a secreted product and exhibited colonization resistance against MRSA in an *in vivo* murine skin model. Using parallel genomic and biochemical investigation, we discovered that *D. incerta* secretes an antimicrobial protein. Sequential protein purification and proteomics analysis identified 24 candidate inhibitory proteins, including a promising peptidoglycan hydrolase candidate. Aided by transcriptional analysis of *D. incerta* and MRSA cocultures, we found that exposure to *D. incerta* leads to decreased MRSA biofilm production. These results emphasize the value in exploring microbial communities across a spectrum of hosts, which can lead to novel therapeutic agents as well as increased understanding of microbial competition.

**IMPORTANCE:** Methicillin-resistant *Staphylococcus aureus* causes significant healthcare burden and can be spread to the human population via livestock transmission. Members of the skin microbiome can prevent MRSA colonization via a poorly-understood phenomenon known as colonization resistance. Here, we studied colonization resistance of *S. aureus* by bacterial inhibitors previously identified from a porcine skin model. We identify a pig skin commensal, *Desemzia incerta*, that reduced MRSA colonization in a murine model. We employ a combination of genomic, proteomic, and transcriptomic analyses to explore the mechanisms of inhibition between *D. incerta* and *S. aureus*. We identify 24 candidate antimicrobial proteins secreted by *D. incerta* that could be responsible for its antimicrobial activity. We also find that exposure to *D. incerta* leads to decreased *S. aureus* biofilm formation. These findings show that the livestock transmission of MRSA can be exploited to uncover novel mechanisms of MRSA colonization resistance.

## INTRODUCTION

*Staphylococcus aureus* is a bacterial pathogen that can infect multiple body sites, including the skin, lung, heart valves, blood, and bone^1^. *S. aureus* is the most commonly identified cause of skin and soft tissue infections (SSTI)^2^ and causes significant healthcare burden^3, 4^; hospitalizations related to *S. aureus* infections in the United States have an estimated annual cost of over 3 billion dollars^5^. Methicillin-resistant *S. aureus* (MRSA) is particularly difficult to treat due to resistance to commonly used antibiotics. Asymptomatic colonization of skin and nares by *S. aureus* is an independent risk factor for subsequent SSTI^6, 7^; colonized individuals are at 4-fold increased risk for *S. aureus* SSTI within 4 years^7^. *S. aureus* spreads among individuals, especially those who are hospitalized^8^ or live in the same household^9^. Livestock workers are also at risk of colonization from livestock-associated MRSA strains^10^. Thus colonization not only increases risk of later *S. aureus* skin infection, but is also a source of community spread of *S. aureus*.

The commensal skin microbiota prevents colonization and invasion by pathogens such as *S. aureus*, a phenomenon known as colonization resistance^11^. One mechanism of colonization resistance is the secretion of antimicrobial peptides and proteins^12^. Host-produced antimicrobial proteins such as LL-37 and defensins may also reduce colonization by *S. aureus*^13, 14^. Both live bacteria and their secreted products can be co-opted for therapeutic effect. For example, *Staphylococcus lugdunesis,* isolated from human nares, produces a cyclic antimicrobial lugdunin. Both *S. lugdunesis* and lugdunin alone can reduce *S. aureus* nasal carriage^15^. Additionally, common skin commensals *S. epidermidis*^16^, *S. hominis*^17^, and *S. capitis*^18^ have all been shown to have antagonistic activity against *S. aureus*. *Cutibacterium acnes* produces a thiopeptide antibiotic, cutimycin, which inhibits staphylococcal species to shape the hair follicle microbiome^19^. Commensal staphylococcal species also produce bacteriocins, antimicrobial peptides which are believed to enable competition in the microbial community while protecting against MRSA and other pathogens^20^. With the looming threat of antimicrobial resistance, harnessing mechanisms of interspecies competition within the skin microbiota could identify novel antimicrobials.

We previously showed that the pig skin microbiota contains a wide phylogenetic range of bacteria that inhibit MRSA in vitro. Not only is pig skin an established in vivo model for human skin ^21^, pigs can become stably colonized with *S. aureus* and MRSA, predominantly livestock-associated strains of MRSA (LA-MRSA)^10^. Livestock colonization is a growing economic and public health concern; livestock workers, particularly swine workers^10^, have high rates of nasal MRSA colonization by LA-MRSA which can lead to subsequent LA-MRSA infections. The most effective method for reducing MRSA colonization in swine herds is eradication of infected herds^22^; however, this creates a large operational and economic burden.

The pig skin microbiome comprises a distinct microbial community from the human skin microbiome. At the genus level, the pig skin microbiome contains *Aerococcus*, *Rothia,* and *Kocuria* species that are not commonly found on human skin. At the species level, the pig skin microbiome contains a distinct set of coagulase-negative *Staphylocci* (CoNS), such as *S. equorum* and *S. haemolyticus*, that are non-overlapping with the CoNS found on human skin^23^. We hypothesized that this distinct microbial community would uncover novel inhibitory mechanisms against *S. aureus*, which may be useful both for therapeutic purposes and for new understanding of interspecies competition in the skin microbiome.

Our previous work identified 37 unique bacterial species that inhibited MRSA in vitro, including 31 species that were not CoNS. We found that three of these pig isolates – *Aerococcus viridans, Rothia aerolata,* and *Desemzia incerta* – cooperatively exhibited colonization resistance against MRSA in a murine skin model^24^. Here we focused on one of these species, *D. incerta*, and investigated more deeply the molecular mechanisms of its interactions with *S. aureus*. We found that *D. incerta* supernatants exhibit MRSA-inhibitory activity *in vitro*. We further found that, at high inoculum, *D. incerta* precolonization exhibits MRSA colonization resistance *in vivo* in a murine skin model. Through a combination of genomic and proteomic approaches, we discovered that *D. incerta* exerts its antimicrobial activity through a secreted antimicrobial protein and identify a peptidoglycan hydrolase protein that is enriched in conditions of high antimicrobial activity. Using transcriptional profiling of *S. aureus* and *D. incerta* cocultures, we discovered that *S. aureus* exposure to *D. incerta* results in significant decrease in *S. aureus* biofilm production. These studies provide greater insight into the microbial and molecular factors underlying resistance to *S. aureus* colonization on skin, and suggest that a peptidoglycan hydrolase might reduce *S. aureus* colonization.

## RESULTS

### A *Desemzia incerta* strain isolated from porcine skin inhibits MRSA via secreted product

We previously cultured 3 bacterial isolates from porcine skin, *A. viridans*, *R. aerolata,* and *D. incerta* that exhibited inhibitory activity against MRSA *in vitro*, and together *in vivo*^24^. To examine the individual mechanisms of inhibition, we tested whether these three isolates secreted antimicrobial molecules. We collected and concentrated cell-free supernatant from bacterial cultures and tested these supernatants for antimicrobial activity using an *in vitro* agar diffusion assay (Figure 1A). Cell-free supernatants were concentrated via a 30 kDa molecular-weight cutoff (MWCO) filtration. Live cell culture or concentrated supernatants were then spotted onto a lawn of MRSA. The presence of a zone of clearing around the spot culture suggested MRSA growth inhibition. Of the three isolates tested, only supernatant from *D. incerta*, a poorly studied Gram-positive rod^25^ (Figure 1B) inhibited MRSA (Figure 1C). This suggests that *D. incerta* secretes an antimicrobial product. We also found that *D. incerta* cells and supernatant inhibited various staphylococcal strains, including human-epidemic MRSA (USA300), livestock-associated MRSA (ST398), and methicillin-sensitive *S. aureus* (strain 502A), as well as the common skin commensal *S. epidermidis.* We also observed inhibition against the Gram-negative skin pathogen *Pseudomonas aeruginosa* (Figure 1C). While the skin pathogen *Streptococcus pyogenes* was inhibited by *D. incerta* live cell cultures, it was not inhibited by *D. incerta* supernatant. Together, these findings suggest that *D. incerta* secretes an antimicrobial product with activity against a range of skin pathogens.

**Figure 1:**
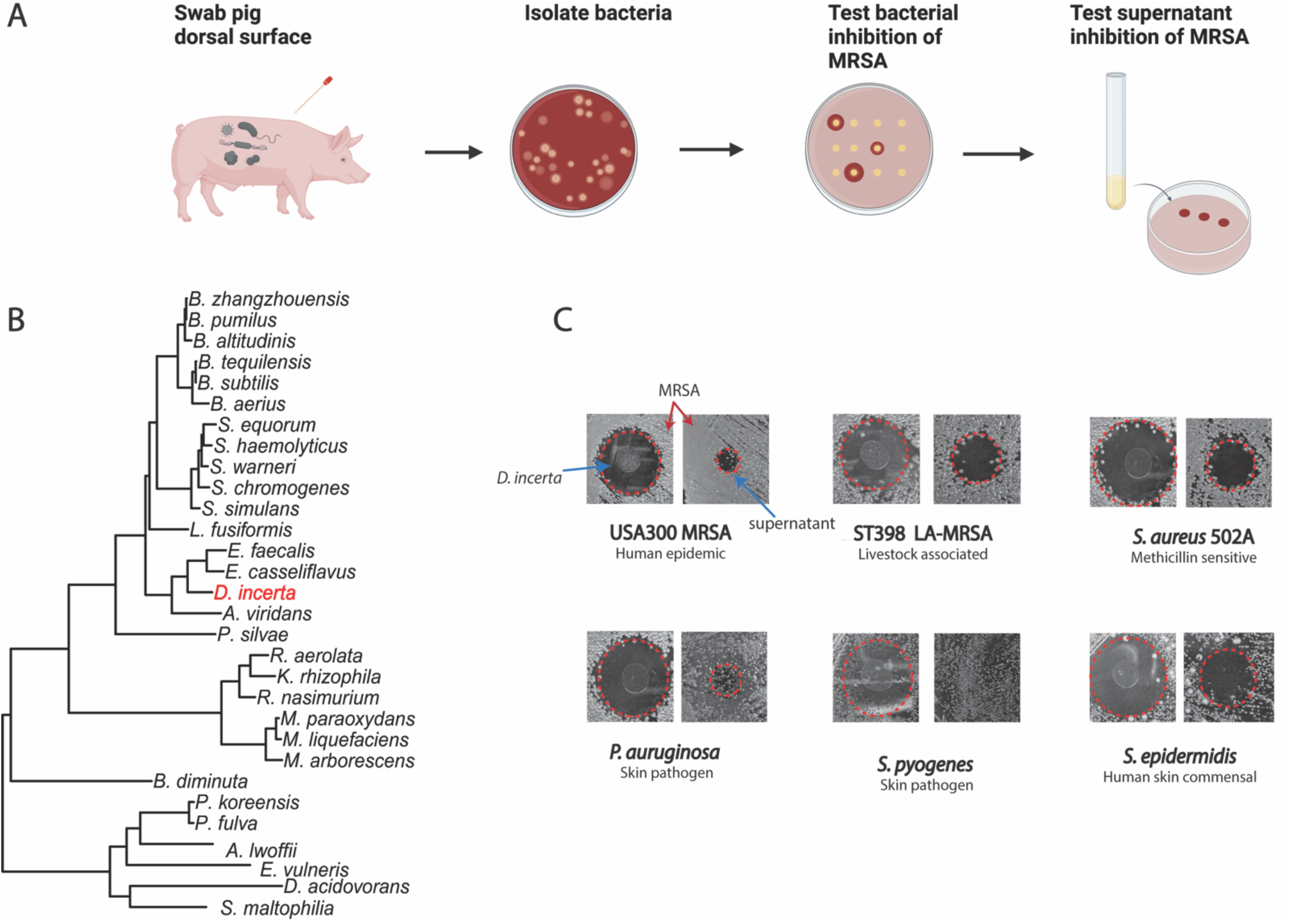
High-throughput screen of pig skin inhibitors reveals *Desemzia incerta* as an inhibitor of MRSA. **A)** Strategy for screening pig skin isolates for inhibitors with secreted MRSA-inhibitory activity. **B)** Dendrogram of unique MRSA-inhibitory bacterial identified from high-throughput screen. Phylogeny was constructed using representative 16S rRNA sequences curated from the Silva database^72^. **C)** Agar diffusion assay of *D. incerta* total cell culture (left) or concentrated supernatant (right) against various target skin pathogens and skin commensals. The edge of the zone of inhibition is demarcated in red dotted lines.

### *D. incerta* decreases MRSA colonization in a murine skin model

We next investigated the ability of *D. incerta* to exert colonization resistance against MRSA in a murine skin colonization model on C57BL/6 mice (Figure 2A). We examined both a pre-colonization (Fig 2B) and de-colonization (Fig 2C) model, where *D. incerta* was either applied prior to (precolonziation) or after (decolonization) MRSA challenge. In both models, 10^9^ CFU *D. incerta* was applied daily for 2 days and 10^8^ CFU USA300 MRSA was applied daily for one day. We found that precolonization with *D. incerta* reduced subsequent MRSA colonization by 60%, but decolonization with *D. incerta* did not have a noticeable effect on MRSA bacterial burden.

**Figure 2:**
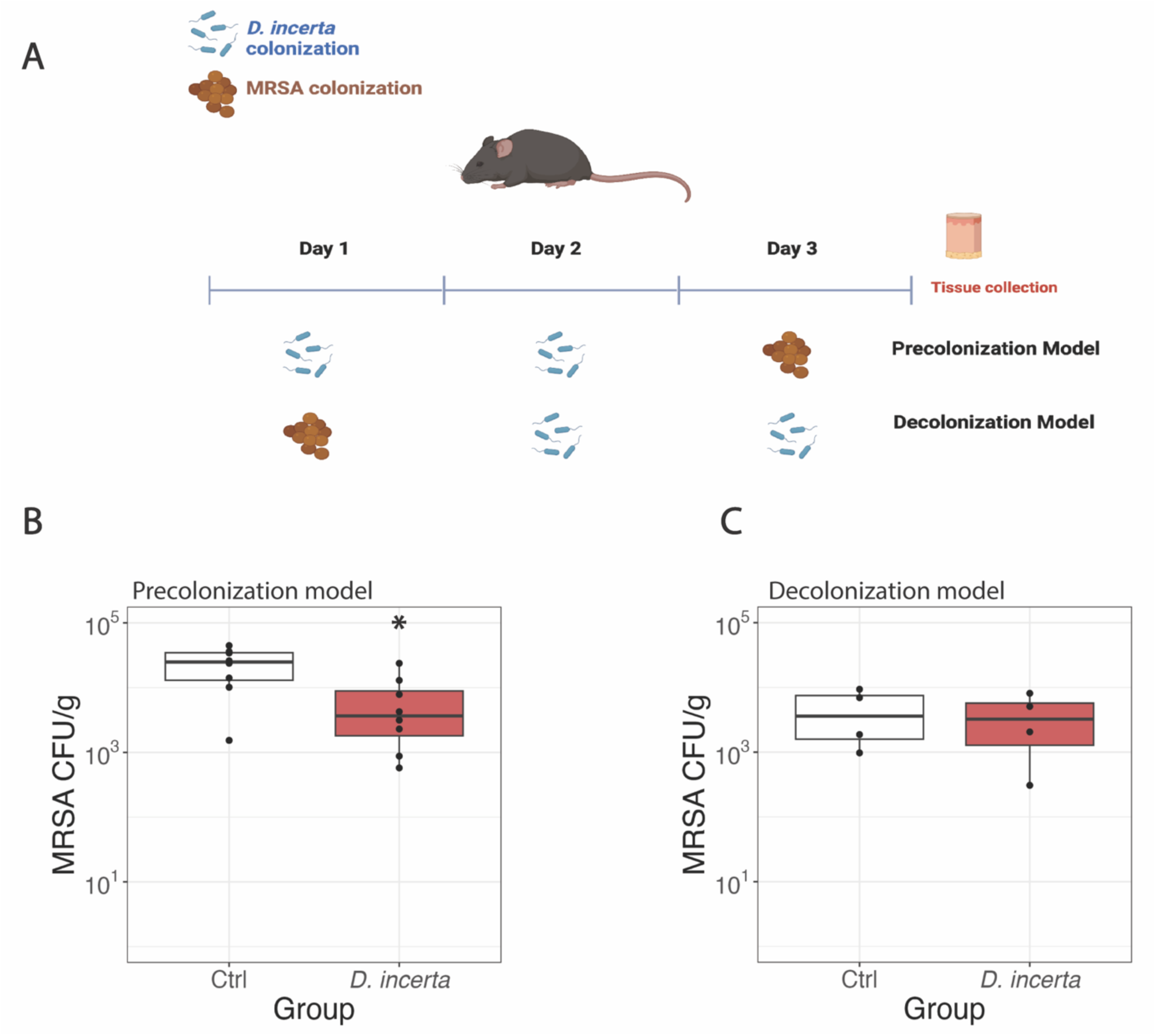
*D. incerta* exhibits colonization resistance against methicillin-resistant *Staphylococcus aureus*. **A)** Experimental design of *in vivo* murine colonization experiments. Mice were shaved and colonized with *D. incerta* for 2 days or MRSA for 1 day in a precolonization or decolonization model. Three 6mm punch biopsies were collected from the mouse dorsal surface and MRSA bacterial load was measured via MRSA-selective CHROMagar. **B)** MRSA bacterial burden after *D. incerta* precolonization. **C)** MRSA bacterial burden after *D. incerta* decolonization. t-test, * p < 0.05

This result differs from our previously reported data, where precolonization by *D. incerta* was not sufficient to reduce MRSA colonization but showed a trend toward statistical significance^24^. In our previous model we used a lower inoculum of *D. incerta* (10^8^ CFU) on a different genetic background (SKH1 hairless mice). Altogether, the data suggests that mono-colonization with *D. incerta* exhibits colonization resistance against MRSA at high innocula, and using a consortia of bacteria as described previously^24^ may increase this anti-MRSA effect.

### Parallel Genomic and Biochemical analyses of *D. incerta* supernatant reveal candidate antimicrobial proteins

To better understand the molecular factors underlying *D. incerta* inhibition of MRSA, we focused on the secreted antimicrobial factor found in *D. incerta* supernatant using a combination of genomic and biochemical approaches. First, we assembled a complete whole genome sequence of *D. incerta* using a hybrid sequencing approach combining Oxford Nanopore long reads and Illumina short reads. This approach allowed us to construct a *de novo* assembly of a complete, circular *D. incerta* genome which included 5 circular plasmids. We used antiSMASH v6^26^ to mine this genome for biosynthetic gene clusters (BGCs) encoding antimicrobial products. antiSMASH employs a homology-based search against a large database of known biosynthetic gene clusters^27^. Fourteen BGCs were identified, all on the bacterial chromosome, and were predicted to encode 9 saccharides, 2 fatty acids, 2 terpenes, and 1 type-3 polyketide (Supplemental Table S1).

To support the genomic analysis, we performed biochemical characterization of the antimicrobial activity present in culture supernatants from *D. incerta*. The antimicrobial activity found in cell-free conditioned media failed to pass through a 50kDa molecular weight cutoff filter and was eliminated after treatment with proteases trypsin or proteinase K (Supplemental Fig S1). These experiments suggested that the antimicrobial activity observed in *D. incerta* supernatant arose from an active protein of >50kDa MW. As our genomic analysis did not predict any BGCs encoding peptide products, it is possible that *D. incerta* produces an antimicrobial protein not encoded by a known BGC.

We next developed a protein purification strategy to isolate the antimicrobial activity from conditioned media (Figure 3A). We combined molecular-weight cutoff concentration with ion-exchange and size-exclusion chromatography to fractionate *D. incerta* cell-free supernatants. The agar diffusion assay described above was used to track activity throughout the purification scheme. After confirming that pooled chromatography products maintained activity against USA300 Rosenbach strain *S. aureus* (Figure 3C), we switched to assaying individual chromatography fractions against *S. epidermidis* (Figure 3B). *S. epidermidis* is a human skin commensal that is safer and more efficient to work with compared to MRSA. At each step, active fractions were pooled and used as the input for subsequent purification steps (Figure 3A). This purification technique yielded 2 active fractions after the final chromatography step (Figure 3B). The combined purification scheme resulted in enrichment for antimicrobial activity; at each step, we noted reduction in total protein concentration coincident with increased or stable antimicrobial activity as measured by agar diffusion assay (Figure 3C).

**Figure 3:**
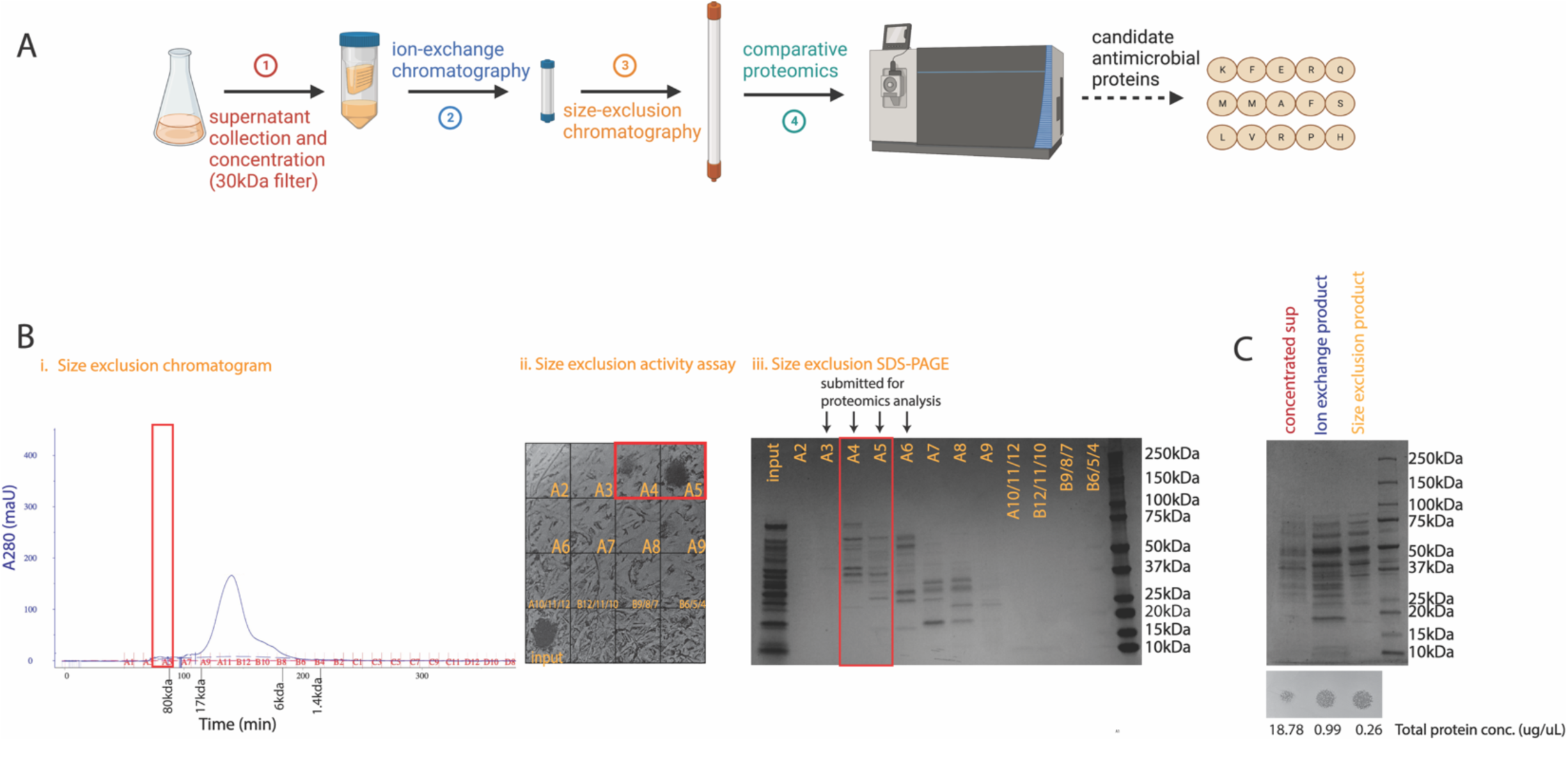
Protein purification and comparative proteomics yields candidate antimicrobial proteins. **A)** Biochemical purification scheme used to generate candidate antimicrobial proteins. 3 biological replicates were submitted for proteomics analysis. **B)** Representative results from the final protein purification step (size exclusion chromatography). One biological replicate is shown. Active fractions, marked in red, are tracked via chromatogram (i), agar diffusion assay against *S. epidermidis* (ii), and SDS-page (iii, 9uL from each sample). Chromatography product from active fractions (A4 and A5) along with neighboring inactive fractions (A3 and A6) were submitted for proteomics analysis. **C)** Representative results across all sequential protein purification steps (molecular weight fractionation, ion exchange chromatography, and size exclusion chromatography). One biological replicate is shown. SDS-PAGE (top, 30μL from each sample), agar diffusion assay against USA300 MRSA (middle) and total protein concentration via nanodrop A280 (bottom) are tracked for pooled active fractions from each purification step.

We submitted the resulting two active fractions from size exclusion chromatography, along with two flanking inactive fractions, for global proteomics analysis to identify proteins that were enriched in the fractions with antimicrobial activity. As *D. incerta* is a poorly studied bacterial species without an empirically validated reference proteome, we constructed a theoretical reference proteome based on predicted open reading frames within the *D. incerta* genome. This approach yielded 24 candidate proteins which were identified as enriched in the active antimicrobial fractions compared to inactive fractions (Table 1). Of particular interest was a putative peptidoglycan hydrolase protein; this class of enzyme can cleave peptidoglycan bonds in Gram-positive and Gram-negative^28^ bacterial cell walls and has been demonstrated before to have antimicrobial activity. For example lysozyme and lysostaphyin, which have both been shown to have anti-staphylococcal activity, are notable members of this family of enzyme^29^. Lysostaphin has also been shown to disrupt *Staphylococcal* biofilms in addition to cleaving peptidoglycan bonds in the cell wall^30^.

**Table 1:**
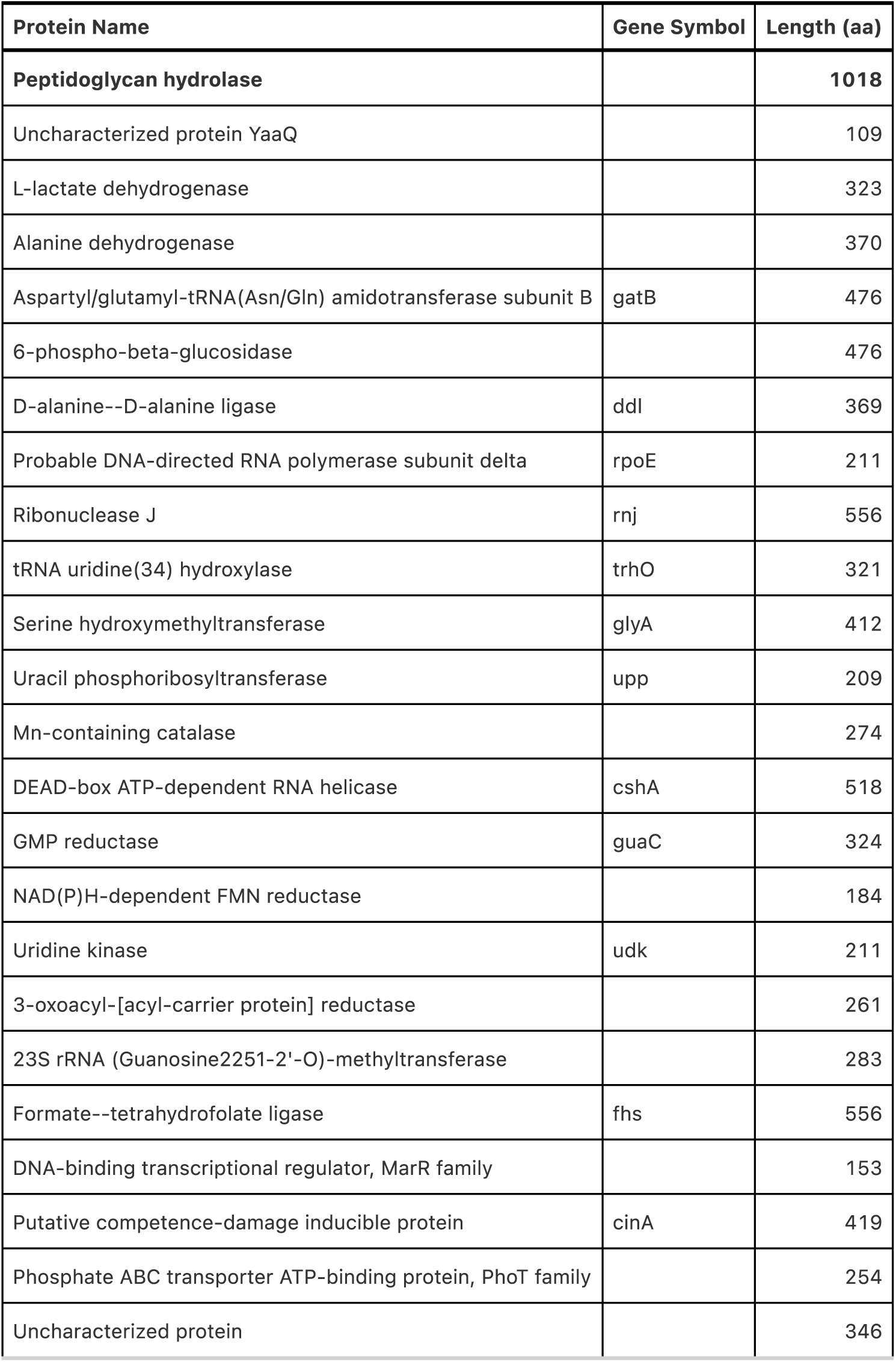
Comparative proteomics reveals candidate antimicrobial proteins. Name, gene symbol, and length in amino acids of 24 proteins found to be enriched in active antimicrobial fractions after protein purification. Ribosomal subunits (e.g. 30S ribosomal subunit) were removed as low-confidence hits. Rows are unsorted.

| Protein Name | Gene Symbol | Length (aa) |
| --- | --- | --- |
| **Peptidoglycan hydrolase** |  | **1018** |
| Uncharacterized protein YaaQ |  | 109 |
| L-lactate dehydrogenase |  | 323 |
| Alanine dehydrogenase |  | 370 |
| Aspartyl/glutamyl-tRNA(Asn/Gln) amidotransferase subunit B | gatB | 476 |
| 6-phospho-beta-glucosidase |  | 476 |
| D-alanine--D-alanine ligase | ddl | 369 |
| Probable DNA-directed RNA polymerase subunit delta | rpoE | 211 |
| Ribonuclease J | rnj | 556 |
| tRNA uridine(34) hydroxylase | trhO | 321 |
| Serine hydroxymethyltransferase | glyA | 412 |
| Uracil phosphoribosyltransferase | upp | 209 |
| Mn-containing catalase |  | 274 |
| DEAD-box ATP-dependent RNA helicase | cshA | 518 |
| GMP reductase | guaC | 324 |
| NAD(P)H-dependent FMN reductase |  | 184 |
| Uridine kinase | udk | 211 |
| 3-oxoacyl-[acyl-carrier protein] reductase |  | 261 |
| 23S rRNA (Guanosine2251-2'-O)-methyltransferase |  | 283 |
| Formate--tetrahydrofolate ligase | fhs | 556 |
| DNA-binding transcriptional regulator, MarR family |  | 153 |
| Putative competence-damage inducible protein | cinA | 419 |
| Phosphate ABC transporter ATP-binding protein, PhoT family |  | 254 |
| Uncharacterized protein |  | 346 |

### Transcriptional response of *S. aureus* to *D. incerta* exposure

In addition to investigating the mechanisms by which *D. incerta* exerts its antimicrobial activity, we also explored the response of *S. aureus* to *D. incerta*. We co-cultured *D. incerta* with either SA113 strain *S aureus* (methicillin-sensitive) or USA300 strain MRSA in a transwell co-culture system (Figure 4A). *D. incerta* liquid cultures were inoculated into the upper transwell and *S. aureus* cultures into the lower transwell at a ratio of 20 *D. incerta* : 1 *S. aureus*. This ratio is comparable to the one used in our *in vitro* agar diffusion assay and our *in vivo* murine colonization studies. The bacterial cultures were separated by a 0.4μm filter, which separates cells but allows for transfer of diffusible molecules through a shared communal media. After 24 hours growth, we measured cell density via OD_600_ and extracted RNA for transcriptomic analysis. We found that the *D. incerta* concentration used in this system resulted in a sub-lethal response by both *S. aureus* strains (Figure 4B, 4C). Exposure to *D. incerta* led to 17% reduction in culture density in *S. aureus* SA113, and no appreciable difference in culture density in USA300 MRSA. *D. incerta* cell density was significantly decreased by >40% in coculture with both *S. aureus* strains, suggesting a co-directional competition between *D. incerta* and *S. aureus*.

**Figure 4:**
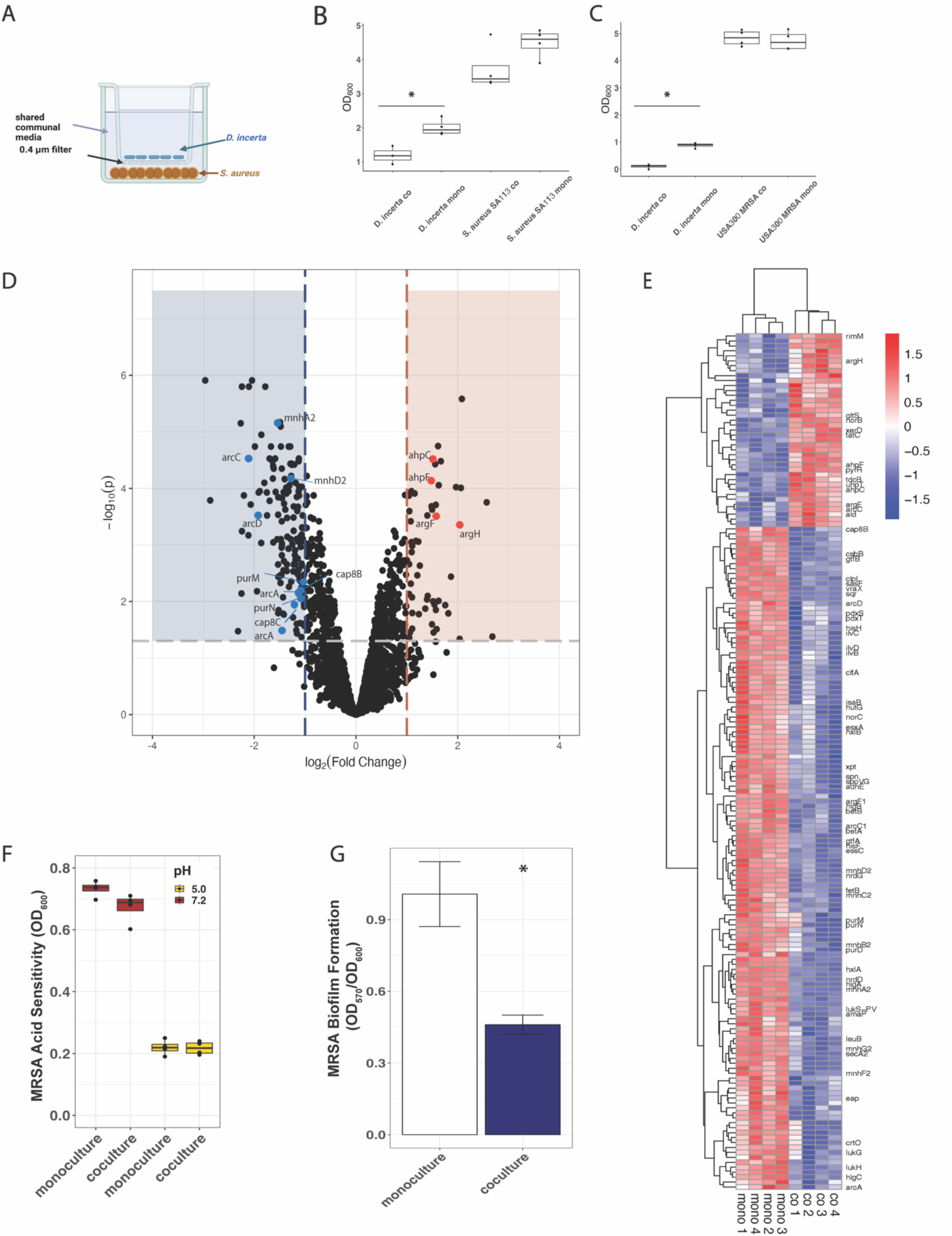
Transcriptional profiling of *S. aureus* reveals changes in biofilm formation after exposure to *D. incerta*. **A)** Transwell coculture experimental design. 4 wells per condition were extracted for RNA-seq analysis. **B)** Bacterial density of *D. incerta* and *S. aureus* SA113 after 24 hrs incubation (OD_600_ of bacterial cultures before RNA extraction) **C)** Bacterial density *D. incerta* and USA300 MRSA after 24 hrs incubation (OD_600_ of bacteria before RNA extraction) **D)** Volcano plot of genes in USA300 MRSA that were differentially expressed during *D. incerta* coculture compared to monoculture. Shaded areas highlight genes whose adjusted p < 0.05 and fold change > 2.0 (blue – downregulated, red – upregulated). Bold points mark genes of interest, which were found within operons containing multiple differentially expressed genes. **E)** Heatmap depicting differentially expressed genes in USA300 MRSA with adjusted p< 0.05 and fold change >2.0. For visualization purposes, genes were ordered by hierarchical clustering based on Pearson correlation. Values were scaled to the row mean. **F)** USA300 MRSA cell density in response to acidified media (pH = 5.0, yellow) compared to neutral pH media (red) and coculture with *D. incerta*. 4 wells per condition were used. **G)** USA300 MRSA biofilm formation measured via crystal violet stain retention (OD_570_) normalized to cell growth (OD_600_). 2 replicates of 3 wells per condition were used. T-test, * p < 0.05

We first examined the transcriptional interaction between *D. incerta* and USA300 MRSA, as this MRSA strain was the focus of our previous *in vitro* and *in vivo* experiments. We focused on the transcriptional response of USA300 MRSA, with particular interest in operons with multiple genes whose transcription was altered in the same direction. 151 genes were significantly downregulated with at least 2-fold change in gene expression (Supplemental Table S2), including multiple genes in the arc operon (*arcA*, *arcC*, *arcD*), the pur operon (*purD*, *purH*, *purM*, *purN*), the mnh family (*mnh*A-G), the luk family (*lukG, lukH, lukS-PV*) and the cap family (*capA*, *cap8B*, *cap8C*) (Figure 4D, Figure 4E). The arc operon encodes a constitutive arginine deaminase system that allows USA300 strain *S. aureus* to survive in acidic microenvironments such as the skin^1^. The pur operon encodes purine biosynthesis^32^; the synthesized purines can then be integrated into downstream nucleic acid synthesis. Mutations in *purM* have been associated with increased susceptibility to environmental stressors such as heat and acidity as well as increased susceptibility to the antibiotic rifampicin^33^. The mnh proteins encode a family of Na^+^/H^+^ transporters that are essential for maintaining cytosol pH and enabling bacterial survival under high salt or high pH conditions^34^.

The luk genes encode a family of *S. aureus* leukocidins, which are important *S. aureus* virulence factors that are well-conserved across *S. aureus* lineages^35^. Leukocidins are thought to protect *S. aureus* from phagocytosis by immune cells. Downregulation of these genes might therefore lead to decreased viability of *S. aureus in vivo*.

Capsular polysaccharide biosynthesis (cap) proteins are required for synthesis of the polysaccharides that form the *S. aureus* capsule. Capsular polysaccharides have been found to protect *S. aureus* from phagocytic killing by human neutrophils^36^. They are believed to be important in cellular adhesion and biofilm formation, though their role in these processes is still unclear: while capsular polysaccharide production is thought to inhibit bacterial adherence and biofilm formation in general, they may be important in maintaining mature biofilms^37^. Defects in capsular polysaccharide synthesis have been associated with decreased nasal colonization of *S. aureus*^38^.

Together, this transcriptional profile of downregulated genes in USA300 *S. aureus* suggest that exposure to *D. incerta* may result in in greater sensitivity to environmental insults, such as acidic pH or high salt concentration, that might be present in the skin microenvironment. Downregulation of *cap* proteins might also lead to changes in biofilm production and decreased ability to colonize skin surfaces, while downregulation of leukocidins might decrease evasion of host immune cells.

In USA300 MRSA, 55 genes were significantly upregulated after coculture with *D. incerta* (Supplemental Table S3). This includes multiple genes in the ahp operon (*ahpC*, *ahpF*) and the arg operon (*argF*, *argH*) (Figure 4D, Figure 4E). ahpFC genes encode alkyl hydroperoxidase, which has been shown to play an important role in bacterial peroxide resistance^39^. *argF* and *argH* are both genes involved in arginine biosynthesis^40^. Notably, the concurrent downregulation of arginine catabolism genes (*arc* operon) and upregulation of arginine biosynthesis genes (*arg* operon) may represent a transcriptional response to low arginine. Taken together, the transcriptional profile of upregulated genes suggests that coculture with *D. incerta* may exert metabolic or oxidative stress onto *S. aureus*.

Though we observed different growth phenotypes between USA300 MRSA and *S. aureus* SA113 after exposure to *D. incerta* (18% growth decrease in SA113 and no growth decrease in USA300 MRSA), we found a remarkably similar transcriptional profile between *S. aureus* SA113 and USA300 MRSA after *D. incerta* exposure (Supplemental Table S4, Supplemental Table S5). Notably, we found a similar downregulation of genes in the arc operon, pur operon, and cap family of genes as well as a similar upregulation of the arg operon.

### Exposure to *D. incerta* decreases *S. aureus* biofilm formation

Because exposure to *D. incerta* led to downregulated *S. aureus* genes involved in pH response and capsule formation, we theorized that exposure to *D. incerta* leads to oxidative or metabolic stress in *S. aureus,* that may hamper the ability of *S. aureus* to survive in acidic conditions or maintain biofilms. We used the same transwell co-culture system as above to test this hypothesis. First, we co-cultured MRSA and *D. incerta* in acidified (pH = 5.0, approximately skin physiologic pH) media compared to standard (pH = 7.2) media. Though acidified media stunted MRSA growth compared to neutral pH media, we did not observe increased acid susceptibility after coculture with *D. incerta* (Figure 4F). Next, we measured MRSA biofilm formation under neutral pH by quantifying retention of crystal violet stain^41^. We noticed a dramatic decrease in MRSA biofilm formation after exposure *D. incerta* (Figure 4G). We also noted a similar response to acidified media and a significant reduction in biofilm formation by *S. aureus* strain SA113 after exposure to *D. incerta* (Supplemental Figure S2).

## DISCUSSION

The phenomenon of colonization resistance is frequently described^42, 43^ but still poorly understood. Better insight into the microbial interactions that shape this phenomenon is important on multiple fronts; first, it can provide greater clarity into the ways that microbial communities behave, which can have important implications for human health^44, 45^. Secondly, competitive microbial interactions can be tapped for their therapeutic potential, enabling discovery of novel antibiotic compounds for which there is urgent need^46^. Here, we focus on a single competitive interaction between the pig skin isolate *D. incerta* and human epidemic strain USA300 MRSA, which we show to exhibit colonization resistance on murine skin. We examined the factors that *D. incerta* uses to initiate this interaction, as well as the response by methicillin-resistant *S. aureus*. We identify 24 candidate antimicrobial proteins secreted by *D. incerta* that might be responsible for its anti-MRSA activity. Of particular interest is a secreted peptidoglycan hydrolase protein, which has been previously proposed to have antimicrobial function. We further profiled the transcriptional response by *S. aureus* and found diminished expression of capsular synthesis genes concurrent with a decrease in biofilm production. These factors might contribute to the colonization resistance phenotype we observe *in vivo*. Notably, *D. incerta* exhibited a different antimicrobial pattern against *S. pyogenes* than against *S. aureus* (Figure 1C), which points to presence of additional antibiotic factors separate from the antimicrobial proteins we focused on here. For example, our genomic analysis indicates that the *D. incerta* genome encodes a type 3 polyketide synthesis (T3PKS) biosynthetic gene cluster (Supplemental Table S1). Polyketides comprise a diverse range of natural products, many of which are antimicrobial^47^. They would also not be expected to pass through the molecular weight cutoff filter we used in our experiments.

While *S. aureus* exposure to *D. incerta* cells and supernatant resulted in clear growth reduction on solid agar diffusion assay, as demonstrated by a zone of clearing, this growth defect was not recapitulated when *S. aureus* was co-cultured with *D. incerta* in liquid culture in our transwell experiments. This difference may be the result of the different nutrient conditions, spatial organization, or culture timing present between the two *in vitro* models we used. The transcriptional profile of *S. aureus* in liquid co-culture pointed to a placation in *S. aureus* virulence through reduced immune evasion (*luk* genes) and decreased capsule synthesis (*cap* genes). Notably these changes were present in both USA300 MRSA and SA113 *S. aureus* despite USA300 MRSA not exhibiting a decrease in cell density, suggesting that this response is not correlated with cell growth. It is therefore unclear whether the reduction in *S. aureus* colonization we observed *in vivo* occurred primarily through reduction in *S. aureus* burden through bacterial antimicrobial factors or through decreased *S. aureus* immune evasion and colonization efficiency.

We focused our studies on a single strain of *S. aureus*, USA300 methicillin-resistant *S. aureus*, because of its high healthcare burden as the leading cause of human epidemic MRSA infections in the United States^48^. Nonetheless, we also observed antimicrobial activity by *D. incerta* against multiple other *S. aureus* strains, including livestock-associated ST398 strain MRSA. Thus, microbiome-derived antimicrobial products might be useful not just for human health but also for reducing the MRSA burden among livestock, which can have both agricultural and public health benefits.

Pigs are an economically important livestock animal that exhibit high rates of MRSA colonization. Pigs and other livestock animals can act as a reservoir for antimicrobial resistance that can cause infection in humans, thus exacerbating an existing public health concern^49^. Here we paradoxically exploited the propensity for MRSA colonization in pigs for therapeutic benefit, by exploring the diverse microbial community on pig skin to identify novel MRSA inhibitors. *D. incerta*, apart from being poorly studied in general, is unique in that it is one of the few MRSA inhibitors isolated from the skin microbiome that is not also a *Staphylococcus* species. A candidate antimicrobial molecule secreted by *D. incerta* that we identified is also much larger (80kDa based on size exclusion chromatography) than other antimicrobial molecules isolated from human skin^15, 16^. The uncovering of *D. incerta*, of which relatively little is known, and its secreted antimicrobial products demonstrates the benefit of exploring alternative microbial communities for new products.

The human microbiome undergoes dynamic exchange with environmental exposures, including other microbial communities. In the context of the skin microbiome, this means that humans may experience transfer of pathogenic bacteria such as *S. aureus* through domestic or occupational exposure, including livestock exposure. This transfer is particularly pronounced between swine, likely due to the similarities between porcine and human skin. Here we show that this shared burden of *S. aureus* colonization can also provide an opportunity for therapeutic discovery and understanding of microbial competition.

## MATERIALS AND METHODS

### *D. incerta* culture and supernatant collection

*D. incerta* from glycerol stocks was streaked onto a blood agar plate and incubated overnight at 37°C. A single large colony was inoculated into 3mL Tryptic Soy Broth (TSB) and incubated overnight at 37°C without shaking. This 3 ml culture was subinoculated into 50mL TSB and incubated overnight at 37°C without shaking. The 50ml culture was subinoculated into 500-1000mL TSB and incubated at room temperature for 3 days without shaking. Cells were pelleted via centrifugation at 15,000g for 15 min and supernatant was filtered through a Stericup .22μm sterile filter. Cell free supernatants were concentrated to <50mL total volume using an Amicon stirred cell model 8050 fitted with a 76mm 30kDa Ultracel MWCO filter (Amicon) or a 76mm 50kDa Biomax MWCO filter (Amicon).

### Agar diffusion assay

*Staphylococcus aureus* was inoculated into 3mL Tryptic Soy Broth (TSB) and incubated at 37°C overnight without shaking. Overnight cultures were diluted to a final OD_600_ of 0.1. 100uL of dilute culture was spread onto a 7cm Tryptic Soy Agar (TSA) plate using glass beads. 5uL of undiluted *D. incerta* culture, concentrated *D. incerta* supernatant, or chromatography product was spotted onto the *S. aureus* lawn. The plate was then incubated overnight at 37°C.

### Construction of phylogenetic tree

Representative 16S ribosomal RNA sequences for each species were curated from the Silva Living Tree Project (LTP) database. For species without a Living Tree Project entry, the longest high-quality sequence from the Reference Non-Redundant (refNR) dataset was used.

Multiple sequence alignment of representative 16S rRNA sequences was generated using MAFFT^50^ v. 7.505 using the L-INS-i setting. A maximum likelihood tree was constructed from the multiple sequence alignment with RAxML^51^ v. 8.2.12 using the best tree from 100 searches. The tree was midpoint rooted in FigTree^52^ v. 1.4.4 and visualized in RStudio.

### Whole genome sequencing and assembly

Genomic DNA was extracted from *D. incerta* cultures using the ZymoResearch QuickDNA Fungal/Bacterial MicroPrep kit. Illumina library preparation and sequencing was performed by the CHOP Microbiome Center, using the Illumina DNA Prep kit and unique dual indexes (Illumina Nextera Index kit) at 1:4 scale reaction volume, and sequenced on the IlluminaHiSeq2500. Trim-galore^53^ v 0.2.4 was used to trim adapter sequences. Quality control of reads was performed before and after read trimming using fastQC^54^ v. 0.11.8 settings. In tandem, high molecular weight genomic DNA was extracted using the NEB Monarch HMW DNA Extraction Kit for Tissue using the standard input workflow for Gram-positive bacteria. For lysis, 80uL of 100mg/mL lysozyme in STET buffer was used, with no other modifications to manufacturer protocol. Library preparation and ONT sequencing of high-molecular weight DNA was performed by SeqCenter. Libraries were prepared using the ONT Genomic DNA by Ligation kit, and sequenced on an ONT MinION with R9 flow cells (R9.4.1). Base calling was performed using Guppy^55^ v 5.0.16 in high-accuracy base calling mode. PoreChop^56^ v 0.2.4 was used to trim adapter sequences. Hybrid assembly using both long and short reads was constructed using Unicycler^55^ v. 0.4.8. Genome annotation was performed by prokka v1.14.6^57^.

### antiSMASH Analysis

The assembled genome fasta was used as input for antiSMASH^58^ v7 in the interactive web browser at (https://antismash.secondarymetabolites.org/) under ‘loose’ settings with all extra features (KnownClusterBlast, ClusterBlast, SubClusterBlast, MiBiG cluster comparison, ActiveSiteFinder, RREFinder, Cluster Pfam analysis, Pfam-based GO annotation, TIGRfam analysis, TFBS analysis) enabled.

### Ion exchange chromatography

Concentrated cell-free supernatant was prepared as described and activity validated against *S. epidermidis* by agar diffusion assay. Bis-tris buffer pH = 5.9 was added to the supernatant to 20mM and loaded using 50mL Superloop onto a 5mL HiTrap QFF column (Cytiva) with gradient elution to 0.5M NaCl in Bis-Tris buffer. 8mL fractions were collected and concentrated using a 30kDA MWCO filter (Amicon). Activity of individual fractions was tested using agar disk diffusion against *S. epidermidis*. Activity of pooled active fractions was validated against USA300 MRSA.

### Size Exclusion Chromatography

Active fractions from the ion-exchange column were pooled, concentrated, and applied to a HiPrep 16/60 S200 column (20mM Bis-Tris pH = 5.9, 50mM NaCl) and 4mL fractions collected. The fractions were concentrated using a 30kDa MWCO filter (Amicon) and activity of individual fractions was tested using agar disk diffusion against *S. epidermidis*. Activity of pooled active fractions was validated against USA300 MRSA.

### Preparation of fractions for mass spectrometry

Fractions from size exclusion chromatography were precipitated by adding NaCl to a final concentration of 100mM, adding four volumes of room temperature acetone, vortexing and incubating at room temperature for 30 minutes. Precipitated proteins were pelleted by spinning at 20k x g for 10 minutes^59^. Pellets were solubilized and digested with the iST kit (PreOmics GmbH, Martinsried, Germany) per manufacturers protocol^60^. Briefly, the resulting pellet was solubilized, reduced and alkylated by addition of SDC buffer containing TCEP and 2-chloroacetamide then heated to 95C for 10minutes. Proteins were enzymatically hydrolyzed for 1.5 hours at 37 °C by addition of LysC and trypsin. The resulting peptides were de-salted, dried by vacuum centrifugation and reconstituted in 0.1% TFA containing iRT peptides (Biognosys Schlieren, Switzerland).

### Mass Spectrometry Data Acquisition

Samples were analyzed on a Q-Exactive HF mass spectrometer (Thermofisher Scientific San Jose, CA) coupled with an Ultimate 3000 nano UPLC system and an EasySpray source. Peptides were loaded onto an Acclaim PepMap 100 75um x 2cm trap column (Thermo) at 5uL/min, and separated by reverse phase (RP)-HPLC on a nanocapillary column, 75 μm id × 50cm 2um PepMap RSLC C18 column (Thermo). Mobile phase A consisted of 0.1% formic acid and mobile phase B of 0.1% formic acid/acetonitrile. Peptides were eluted into the mass spectrometer at 300 nL/min with each RP-LC run comprising a 90 minute gradient from 3% B to 45% B.

The mass spectrometer was set to repetitively scan m/z from 300 to 1400 (R = 240,000) followed by data-dependent MS/MS scans on the twenty most abundant ions, minimum AGC 1e4, dynamic exclusion with a repeat count of 1, repeat duration of 30s, and resolution of 15000. The AGC target value was 3e6 and 1e5, for full and MSn scans, respectively. MSn injection time was 160 ms. Rejection of unassigned and 1+,6-8 charge states was set.

### System Suitability and Quality Control

The suitability of Q Exactive HF instrument was monitored using QuiC software (Biognosys, Schlieren, Switzerland) for the analysis of the spiked-in iRT peptides. Meanwhile, as a measure for quality control, we injected standard *E. coli* protein digest prior to and after injecting sample set, and collected the data in the Data Dependent Acquisition (DDA) mode. The collected data were analyzed in MaxQuant^61^ and the output was subsequently visualized using the PTXQC^62^ package to track the quality of the instrumentation.

### MS data processing and analysis

The raw files for DDA analysis were processed with FragPipe^63^ version 20.0 using its Default workflow. The reference *Desemzia incerta* proteome from UniProt (2,143 canonical and isoform proteins) was concatenated with the self-generated reference proteome and common protein contaminants and used for the search. The default MS1 quantification was performed using IonQuant algorithm through FragPipe without match between runs option. Data processing and statistical analysis was performed in R. The raw MS1 intensity data were log2 transformed and normalized by subtracting the median for each sample. After data filtering, Limma t-test was employed to identify differentially abundant proteins between high and no-activity fractions using P.Value < 0.05 as significant threshold. The proteins that were exclusively detected in one experimental group were also reported for further bioinformatics analysis.

### Transwell co-cultures

*S. aureus* and *D. incerta* overnight cultures were grown as described. *S. aureus* was diluted to a final OD_600_ of 0.05. *D. incerta* cultures were concentrated to a final OD_600_ of 2.0. 1mL of dilute *S. aureus* culture or TSB control was loaded into the bottom well of a Corning Transwell 12-well cell culture plate (0.4um pore size). 500uL of *D. incerta* culture or TSB control was loaded into the top well of the corresponding transwell insert. Plates were incubated at 37°C overnight without shaking. The contents of the bottom and top well were collected separately into 1.5 mL microcentrifuge tubes. 100uL was reserved for OD_600_ measurements (Model name, Eppendorf). The remainder was centrifuged at >10,000g for 1 min. The supernatant was decanted and the cell pellet was stored at -80°C.

### Growth in Acidified Media

Tryptic Soy Broth was prepared according to manufacturer specifications and found to have pH = 7.2. No modifications were made to this standard TSB in the neutral pH condition. To create acidic media, concentrated acetic acid was added dropwise to standard TSB until the pH reached 5.0. *D. incerta* and *S. aureus* were seeded into transwell culture plates at the ratios described above in either standard media or acidic media and incubated overnight at 37°C. The top well containing *D. incerta* was removed and OD_600_ of the *S. aureus* culture was measured on a BioTek Synergy HT microplate reader.

### Biofilm Formation Assay

*D. incerta* and *S. aureus* were seeded into transwell tissue culture plates as described above into TSB media containing 0.5% dextrose. Cultures were incubated for 24 hours at 37°C. The top well containing *D. incerta* was discarded and *S. aureus* cultures in the bottom well were gently resuspended. OD_600_ of the *S. aureus* was measured on a microplate reader and the *S. aureus* cultures discarded. The plate was rinsed vigorously in deionized water and the wells were stained with 500μL crystal violet solution. After 10 minutes, the crystal violet stain was removed and the plate was rinsed vigorously in deionized water and allowed to dry for 5 minutes. The remaining crystal violet stain was solubilized in 500μL 30% acetic acid, which was diluted in equal volume PBS. OD_570_ of the solubilized crystal violet was measured on a microplate reader.

### RNA Extraction and sequencing

RNA from frozen bacterial cell pellets was extracted using the Zymo Research Direct-zol RNA purification kit. RNA-sequencing was performed by SeqCenter using their standard RNA sequencing methods. In brief, samples were DNAse treated with Invitrogen DNAse (RNAse free). Library preparation was performed using Illumina’s Stranded Total RNA Prep Ligation with Ribo-Zero Plus kit and 10bp IDT for Illumina indices. Sequencing was done on a NextSeq2000 giving 2x51 bp reads. Demultiplexing, quality control, and adapter trimming was performed with bcl-convert (v3.9.3).

### RNA sequencing analysis

Quality control of RNA reads was performed using fastqc^54^ v 0.11.9. Reads were mapped to coding sequences from the *S. aureus* representative genome <u>NC_010079.1</u> using Kallisto^64^ v 0.48.0. Mapped genes were filtered and normalized using edgeR^65^ v 3.40.2 and limma^66^ v 3.54.2 packages in Rstudio v 4.2.2. GO terms were assigned using the Uniprot^67^ IDmapping tool and GO enrichment was performed using the goseq^68^ v 1.50.0 package in Rstudio^69^.

### Mouse Colonization Resistance Experiments

All mouse procedures were performed under protocols approved by the University of Pennsylvania IACUC. Seven week old female C57BL/6 mice were allowed to acclimate for 1 week before experimentation began. Mice were given ad libitum access to food and water. *D. incerta* was grown in 3mL liquid TSB overnight at 37°C without shaking, then subinoculated into 50mL TSB overnight at 37°C without shaking. MRSA isolates were grown overnight in TSB media at 200rpm. On the following day, OD_600_ measurement was used to standardize inoculums, and pellets resuspended in TSB to acquire 1x10^9^ CFU/ml inoculums (1x10^8^ for MRSA). Mice were shaved across their entire back and cage changed one day prior to initial colonization. No additional cage changes were performed throughout the duration of the experiment. Inocula of MRSA (10^8^) or *D. incerta* (10^9^) were prepared from overnight cultures resuspended in 100μL TSB media using the conversion 1OD = 3.3e8 CFU/mL for *S. aureus* and 1OD = 3.5e8CFU/ml for *D. incerta*. Mice were anesthetized and inocula were pipetted onto their shaved backs and spread with a foam-tipped applicator. This procedure was repeated for 3 days with experimental groups as described. 24 hours after the final inoculum was applied, mice were euthanized. 3x 6mm skin punch biopsies were collected into a pre-weighted Eppendorf tube containing 500ul TSB and a ¼” ceramic bead (MP biomedical). Samples were weighed and agitated for 30 min on a VortexGenie 2 with horizontal tube attachment. 100uL of liquid was spread onto a blood agar or MRSA selective CHROMagar (*S. aureus* CHROMagar^TM^ + 5mg/mL cefoxitin) and incubated overnight at 37°C.

### Microbial Strains

Strains will be made available upon request.

| Strain | Source |
| --- | --- |
| *Staphylococcus aureus* subsp. aureus Rosenbach. StrainTCH1516 [USA300-HOU-MR] | ATCC cat.# BAA-1717 |
| *Staphylococcus aureus* 502A subsp. aureus Rosenbach | ATCC # 27217 serologic type (b)c1 |
| *Staphylococcus aureus* S0385 (LA-MRSA) | BEI Resourcess (ATCC) Cat# NR-28983 |
| *Streptococcus pyogenes* (Strain Designation: SF370; M1 GAS ) | ATCC Lot # 70012149 |
| *Staphylococcus epidermidis* (EGM 2-01) | Isolated from healthy human skin<br>Initially described in Uberoi *et al.*<sup>70</sup> |
| *Pseudomonas aeruginosa* (DOERN #1896) | Isolated from diabetic foot ulcer<br>Initially described in Gardner *et al.*<sup>71</sup> |
| *Aerococcus viridans* (EGM 1-16) | Isolated from porcine skin<br>Initially described in Wei *et al.*<sup>24</sup> |
| *Rothia aerolata* (EGM 1-37) | Isolated from porcine skin<br>Initially described in Wei *et al.*<sup>24</sup> |
| *Desemzia incerta* (EGM 1-39) | Isolated from porcine skin<br>Initially described in Wei *et al.*<sup>24</sup> |

### Data Availability

Genome sequences are available in NCBI databases under BioProject PRJNA974032, Biosample SAMN35158552, GenBank CP CP126128.1 (chromosome), NZ_CP126129.1 (plasmid 1), NZ_CP126130.1 (plasmid 2), NZ_CP126130.1 (plasmid 3), NZ_CP126132.1 (plasmid 4), NZ_CP126133.1 (plasmid 5), SRA SRR24674626 (long read) and SRR24710524 (short read).

RNAseq data is available in the Gene Expression Omnibus (GEO) under series GSE239513.

## ACKNOWLEDGEMENTS

We thank the veterinary staff, residents, and faculty of the University Lab Animal Resources (ULAR) for their care for the animals. We thank current and former members of the Grice lab and the Department of Dermatology for critical discussion and review of the work. This work was funded by the following grants to EAG from the NIH, NIAMS (R01AR006663), NINR (R01NR015639), the Burroughs Wellcome Fund PATH Award, the University of Pennsylvania Linda Pechenik Montague Investigator Award, and the Dermatology Foundation Sun Pharma Research Award. This research was also supported by the Penn Skin Biology and Disease Resource-based Center (Penn SBDRC supported by NIH/NIAMS P30AR069589). LF was supported by the Penn Dermatology Research T32 Training Grant (NIH/NIAMS T32AR007465); MW was supported by the Bacterial Pathogenesis T32 Training Grant, (NIH/NIAID 5T32AI141393).

## SUPPLEMENTAL MATERIALS

**Table S1:** Summary of antiSMASH analysis of *D. incerta* complete genome. ‘Loose’ setting was used with all additional features enabled.

| Class | Length | Most Similar Known Cluster (Class) | Similarity (Gene Content) | Number of Genes |
| --- | --- | --- | --- | --- |
| saccharide | 24419 |  |  | 22 |
| saccharide | 30014 |  |  | 26 |
| terpene | 20142 |  |  | 18 |
| saccharide | 30822 |  |  | 28 |
| saccharide | 39794 | oryzanaphthopyran (polyketide) | 6% | 36 |
| saccharide | 21203 |  |  | 24 |
| T3PKS | 41196 | kijanimicin (polyketide) | 4% | 40 |
| saccharide | 38793 |  |  | 37 |
| saccharide | 22906 |  |  | 18 |
| terpene, saccharide | 32766 |  |  | 28 |
| saccharide | 39556 |  |  | 39 |
| fatty acid | 24536 | xantholipin (polyketide) | 4% | 29 |
| saccharide | 30988 |  |  | 26 |
| saccharide, fatty acid | 103495 | capsular polysaccharide (saccharide:<br>exopolysaccharide) | 13% | 71 |

**Figure S1:**
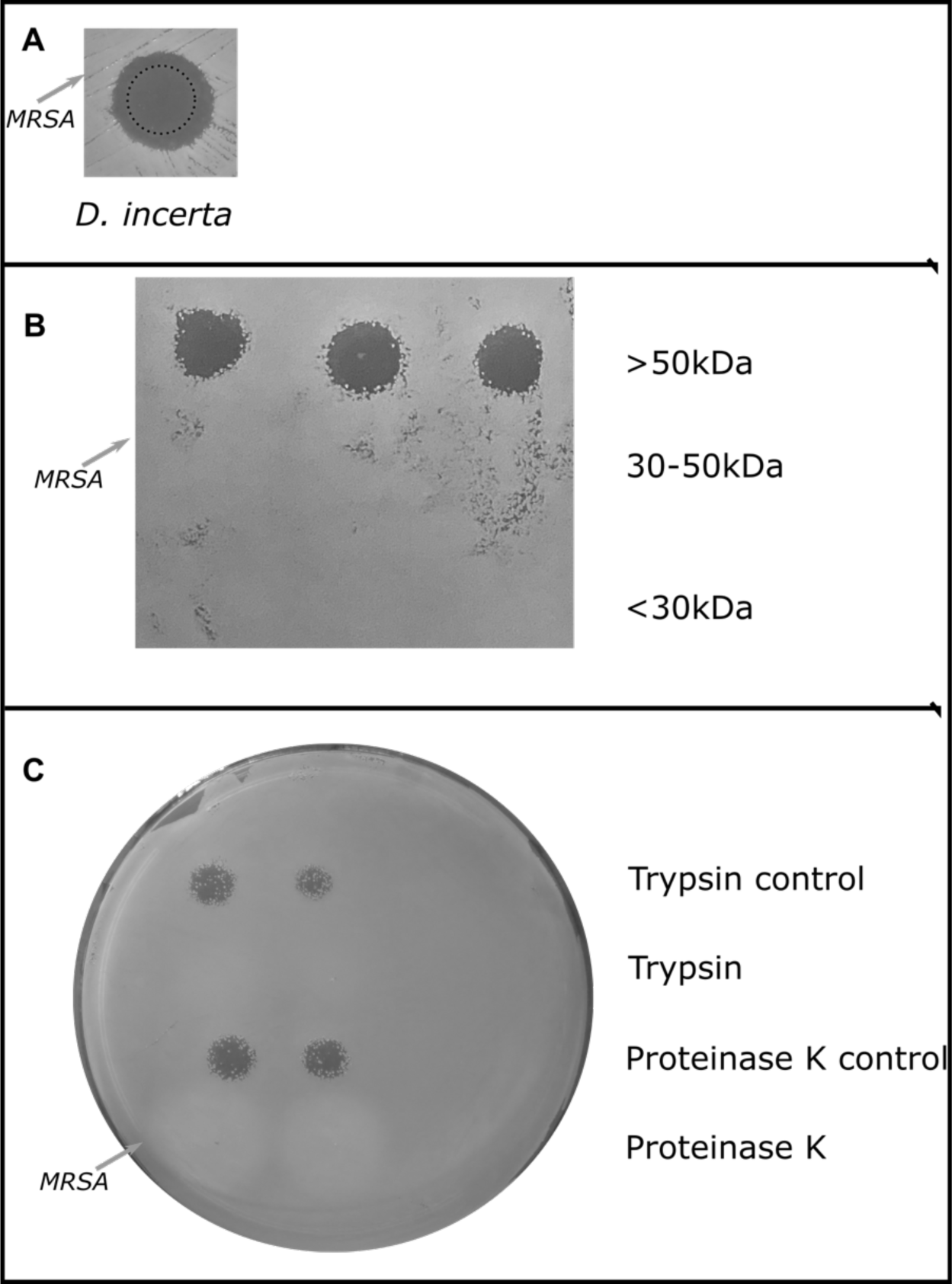
*D. incerta* supernatant is >50kDa and sensitive to protease digestion. **A)** 5 μL *D. incerta* cell culture (OD ∼ 1.0) was spotted onto a lawn of USA300 MRSA (100 μL, OD = 0.1) on Trypic Soy Agar (TSA) and incubated overnight at 37°C. **B)** Conditioned media was collected from *D. incerta* overnight cell cultures and concentrated 20x using a 50kDa molecular weight cutoff (MWCO) filter. The retentate was kept as >50kDa fraction, while the flow through was kept as <50kDa fraction. The flow through was then concentrated 20x in a 30kDa MWCO filter. The retentate was kept as >30kDa but <50kDa fraction, and the flow through was kept as <30kDa fraction. Fractions were spotted on a lawn of MRSA as in (A). Rows represent technical triplicates. **C)** Concentrated supernatant was collected as described in (B). Concentrated supernatant with confirmed antimicrobial activity was incubated with either 60 BAEE units Trypsin, 20 mAU Proteinase K, or the corresponding buffer control for 4 hrs at 37°C. Protease treated samples were tested for MRSA inhibition via the agar diffusion assay described in A. Rows represent technical duplicates.

**Table S2:** USA300 MRSA downregulated genes after exposure to *D. incerta*. Genes with adjusted p val < 0.05 and fold-change >2 are included. Rows are sorted by fold change.

| RefSeqID | log<sub>2</sub>(FC) | adj. p val | Gene Symbol | Protein |
| --- | --- | --- | --- | --- |
| WP_000594519.1 | -2.96 | 1.2E-06 | hlgA | bi-component gamma-hemolysin HlgAB subunit A |
| WP_000669519.1 | -2.87 | 1.6E-04 | spn | myeloperoxidase inhibitor SPIN |
| WP_000595263.1 | -2.32 | 3.3E-02 |  | transcriptional activator RinB |
| WP_000601724.1 | -2.26 | 7.1E-06 | nrdG | anaerobic ribonucleoside-triphosphate reductase activating protein |
| WP_001549197.1 | -2.25 | 7.2E-03 |  | delta-lysin family phenol-soluble modulin |
| WP_000793596.1 | -2.24 | 1.6E-06 | argF | ornithine carbamoyltransferase |
| WP_025175201.1 | -2.24 | 5.7E-04 |  | hypothetical protein |
| WP_000730043.1 | -2.20 | 1.3E-04 |  | SDR family oxidoreductase |
| WP_000036757.1 | -2.12 | 3.0E-05 | arcC | carbamate kinase |
| WP_000475805.1 | -2.11 | 6.8E-04 |  | universal stress protein |
| WP_001071717.1 | -2.10 | 1.6E-06 | nrdD | anaerobic ribonucleoside-triphosphate reductase |
| WP_001093553.1 | -2.10 | 3.0E-05 |  | DM13 domain-containing protein |
| WP_001005511.1 | -2.07 | 1.9E-04 |  | fructosamine kinase family protein |
| WP_000777188.1 | -2.04 | 1.2E-06 |  | metal-dependent hydrolase |
| WP_001549607.1 | -1.99 | 1.8E-05 |  | MAP domain-containing protein |
| WP_000272781.1 | -1.94 | 6.6E-03 |  | Crp/Fnr family transcriptional regulator |
| WP_000239610.1 | -1.93 | 3.0E-04 |  | phospholipase |
| WP_001237629.1 | -1.91 | 1.2E-04 |  | hypothetical protein |
| WP_000791407.1 | -1.89 | 1.1E-04 | lukH | bi-component leukocidin LukGH subunit H |
| WP_000595324.1 | -1.86 | 9.2E-04 | lukG | bi-component leukocidin LukGH subunit G |
| WP_000752917.1 | -1.86 | 1.1E-05 |  | CsbD family protein |
| WP_001056223.1 | -1.80 | 8.0E-05 | clfA | MSCRAMM family adhesin clumping factor ClfA |
| WP_000206639.1 | -1.78 | 1.6E-06 |  | ArgE/DapE family deacylase |
| WP_000916713.1 | -1.75 | 1.2E-04 | hlgC | bi-component gamma-hemolysin HlgCB subunit C |
| WP_000344142.1 | -1.75 | 3.0E-04 | arcD | arginine-ornithine antiporter |
| WP_001789452.1 | -1.70 | 3.0E-05 |  | acyl CoA:acetate/3-ketoacid CoA transferase |
| WP_000558975.1 | -1.70 | 1.7E-04 |  | alpha/beta hydrolase |
| WP_001177186.1 | -1.67 | 1.8E-05 |  | hypothetical protein |
| WP_000066521.1 | -1.64 | 3.3E-05 | betA | choline dehydrogenase |
| WP_000036327.1 | -1.63 | 6.7E-05 |  | DUF1648 domain-containing protein |
| USA300HOU_RS16045 | -1.63 | 4.0E-05 | hisF | imidazole glycerol phosphate synthase subunit HisF |
| WP_000857159.1 | -1.62 | 9.0E-04 |  | zinc-dependent alcohol dehydrogenase family protein |
| WP_000783428.1 | -1.62 | 4.5E-05 | hlgB | bi-component gamma-hemolysin HlgAB/HlgCB subunit B |
| WP_000071901.1 | -1.59 | 3.0E-05 |  | pyridoxamine 5'-phosphate oxidase family protein |
| WP_001077722.1 | -1.58 | 1.2E-04 |  | YjiH family protein |
| WP_000708242.1 | -1.55 | 7.9E-04 |  | antibiotic biosynthesis monooxygenase |
| WP_001096791.1 | -1.53 | 2.2E-03 |  | lantibiotic protection ABC transporter ATP-binding subunit |
| WP_001008401.1 | -1.53 | 2.9E-04 |  | class I adenylate-forming enzyme family protein |
| WP_000446299.1 | -1.53 | 1.6E-02 | capA | capsular polysaccharide type 5/8 biosynthesis protein CapA |
| WP_001548082.1 | -1.53 | 1.4E-04 |  | thermonuclease family protein |
| WP_000196911.1 | -1.53 | 3.7E-03 |  | ACT domain-containing protein |
| WP_000060776.1 | -1.53 | 6.8E-06 | mnhA2 | Na<sup>+</sup>/H<sup>+</sup> antiporter Mnh2 subunit A |
| WP_000368687.1 | -1.52 | 1.4E-02 |  | hypothetical protein |
| WP_000422858.1 | -1.49 | 6.8E-06 | hxlA | 3-hexulose-6-phosphate synthase |
| WP_000635623.1 | -1.48 | 4.7E-03 | hisH | imidazole glycerol phosphate synthase subunit HisH |
| WP_000587958.1 | -1.48 | 8.0E-06 | vraX | C1q-binding complement inhibitor VraX |
| WP_000142189.1 | -1.48 | 3.4E-03 |  | acyl-CoA dehydrogenase family protein |
| WP_000277968.1 | -1.47 | 7.9E-04 | hutG | formimidoylglutamase |
| WP_000680947.1 | -1.47 | 1.8E-05 | secA2 | accessory Sec system translocase SecA2 |
| WP_000581792.1 | -1.47 | 3.4E-03 |  | heavy metal-associated domain-containing protein |
| WP_000810443.1 | -1.46 | 1.1E-03 |  | hypothetical protein |
| WP_001075826.1 | -1.46 | 3.8E-04 |  | M20 family metallopeptidase |
| WP_000129411.1 | -1.46 | 3.2E-02 | arcA | arginine deiminase |
| WP_000735558.1 | -1.46 | 4.5E-05 |  | hypothetical protein |
| WP_001044915.1 | -1.46 | 1.5E-03 |  | tyrosine-type recombinase/integrase |
| WP_000586140.1 | -1.45 | 1.0E-03 |  | hypothetical protein |
| WP_000448901.1 | -1.45 | 9.4E-04 |  | alpha/beta hydrolase |
| WP_001255791.1 | -1.45 | 1.5E-04 | ilvD | dihydroxy-acid dehydratase |
| WP_000154514.1 | -1.43 | 8.4E-03 |  | 3-hydroxyacyl-CoA dehydrogenase/enoyl-CoA hydratase family protein |
| WP_000571736.1 | -1.43 | 1.6E-02 | hisA | 1-(5-phosphoribosyl)-5-((5-phosphoribosylamino)methylideneamino)imidazole-4-carboxamide isomerase |
| WP_000145163.1 | -1.42 | 2.8E-03 |  | sterile alpha motif-like domain-containing protein |
| WP_000163283.1 | -1.42 | 1.2E-04 |  | type 1 glutamine amidotransferase domain-containing protein |
| WP_000410771.1 | -1.42 | 1.7E-02 |  | HdeD family acid-resistance protein |
| WP_001051594.1 | -1.42 | 1.9E-03 | hxlB | 6-phospho-3-hexuloisomerase |
| WP_000763768.1 | -1.41 | 1.9E-04 |  | general stress protein |
| WP_000421701.1 | -1.40 | 6.9E-05 | betB | betaine-aldehyde dehydrogenase |
| WP_000955777.1 | -1.39 | 2.4E-04 | adhE | bifunctional acetaldehyde-CoA/alcohol dehydrogenase |
| WP_000762708.1 | -1.37 | 9.7E-05 |  | putative metal homeostasis protein |
| WP_001037807.1 | -1.37 | 1.2E-04 |  | DoxX family protein |
| WP_001795496.1 | -1.36 | 2.9E-03 | crtO | glycosyl-4,4'-diaponeurosporenoate acyltransferase |
| WP_000655692.1 | -1.36 | 2.2E-03 |  | aminotransferase class I/II-fold pyridoxal phosphate-dependent enzyme |
| WP_001013484.1 | -1.35 | 3.3E-04 |  | cation diffusion facilitator family transporter |
| WP_000581552.1 | -1.35 | 7.2E-04 |  | lantibiotic immunity ABC transporter MutE/EpiE family permease subunit |
| USA300HOU_RS10525 | -1.33 | 1.2E-03 | eap | extracellular adherence protein Eap/Map |
| WP_000827000.1 | -1.32 | 1.1E-04 |  | hypothetical protein |
| USA300HOU_RS15940 | -1.32 | 1.8E-05 | sqr | type II sulfide:quinone oxidoreductase Sqr |
| WP_000030057.1 | -1.32 | 5.5E-03 |  | aspartate aminotransferase family protein |
| WP_000072147.1 | -1.31 | 8.8E-05 | fetB | iron export ABC transporter permease subunit FetB |
| WP_000277602.1 | -1.29 | 1.8E-05 |  | universal stress protein |
| WP_001058981.1 | -1.29 | 3.7E-05 | clpL | ATP-dependent Clp protease ATP-binding subunit ClpL |
| WP_000214557.1 | -1.28 | 7.9E-04 | ilvC | ketol-acid reductoisomerase |
| WP_000158433.1 | -1.28 | 6.3E-05 | gtfA | accessory Sec system glycosyltransferase GtfA |
| WP_000950548.1 | -1.27 | 6.0E-05 | mnhD2 | Na<sup>+</sup>/H<sup>+</sup> antiporter Mnh2 subunit D |
| WP_000690439.1 | -1.27 | 4.4E-03 | pdxT | pyridoxal 5'-phosphate synthase glutaminase subunit PdxT |
| WP_000215236.1 | -1.27 | 3.7E-05 |  | Asp23/Gls24 family envelope stress response protein |
| WP_000825819.1 | -1.26 | 7.4E-05 | amaP | alkaline shock response membrane anchor protein AmaP |
| WP_000593153.1 | -1.26 | 3.8E-05 | gtfB | accessory Sec system glycosylation chaperone GtfB |
| WP_000094576.1 | -1.25 | 9.8E-04 |  | 2-isopropylmalate synthase |
| WP_001154618.1 | -1.24 | 1.9E-02 |  | thiolase family protein |
| WP_000034728.1 | -1.23 | 2.9E-03 | pdxS | pyridoxal 5'-phosphate synthase lyase subunit PdxS |
| WP_000793009.1 | -1.23 | 3.3E-04 |  | nucleobase:cation symporter-2 family protein |
| WP_001206093.1 | -1.22 | 5.6E-04 |  | aldehyde dehydrogenase family protein |
| WP_000616642.1 | -1.22 | 2.1E-04 | mnhF2 | Na<sup>+</sup>/H<sup>+</sup> antiporter Mnh2 subunit F |
| WP_000047820.1 | -1.22 | 2.2E-03 | ilvB | biosynthetic-type acetolactate synthase large subunit |
| WP_000702782.1 | -1.21 | 3.8E-05 | csbB | lipoteichoic acid-specific glycosylation protein CsbB |
| WP_000565302.1 | -1.21 | 1.2E-02 | cap8C | type 8 capsular polysaccharide synthesis protein Cap8C |
| WP_000183771.1 | -1.21 | 1.1E-02 |  | (S)-acetoin forming diacetyl reductase |
| WP_000175846.1 | -1.20 | 1.5E-03 |  | PLP-dependent aspartate aminotransferase family protein |
| WP_000661906.1 | -1.20 | 1.2E-03 | mnhB2 | Na<sup>+</sup>/H<sup>+</sup> antiporter Mnh2 subunit B |
| WP_001048985.1 | -1.19 | 2.2E-03 | mnhC2 | Na<sup>+</sup>/H<sup>+</sup> antiporter Mnh2 subunit C |
| WP_000910581.1 | -1.19 | 3.0E-03 |  | hypothetical protein |
| WP_001108126.1 | -1.19 | 1.9E-03 |  | sucrose-specific PTS transporter subunit IIBC |
| WP_000406611.1 | -1.18 | 8.0E-05 | mnhG2 | Na<sup>+</sup>/H<sup>+</sup> antiporter Mnh2 subunit G |
| WP_000052078.1 | -1.17 | 3.2E-02 |  | hypothetical protein |
| WP_000174578.1 | -1.17 | 7.9E-04 |  | DNA-binding protein |
| WP_000136166.1 | -1.16 | 2.5E-02 | argF | ornithine carbamoyltransferase |
| WP_000769720.1 | -1.16 | 1.4E-04 |  | PepSY domain-containing protein |
| WP_001071973.1 | -1.16 | 3.7E-02 | mnhE2 | Na<sup>+</sup>/H<sup>+</sup> antiporter Mnh2 subunit E |
| WP_000002683.1 | -1.15 | 2.5E-04 |  | DUF2273 domain-containing protein |
| WP_001024094.1 | -1.14 | 1.7E-03 |  | SDR family oxidoreductase |
| WP_000356963.1 | -1.14 | 3.3E-03 |  | hypothetical protein |
| WP_000655875.1 | -1.14 | 7.0E-03 |  | DUF5081 family protein |
| WP_000160456.1 | -1.14 | 2.1E-04 |  | NAD(P)/FAD-dependent oxidoreductase |
| WP_001060912.1 | -1.13 | 8.7E-04 |  | aldehyde reductase |
| WP_000931237.1 | -1.13 | 3.3E-05 |  | aldo/keto reductase |
| WP_000691766.1 | -1.13 | 1.1E-04 |  | RidA family protein |
| WP_001044560.1 | -1.12 | 2.7E-03 | isaB | immunodominant staphylococcal antigen IsaB |
| WP_001151900.1 | -1.12 | 3.0E-05 | sasF | cell-wall-anchored protein SasF |
| WP_000769689.1 | -1.12 | 1.3E-04 |  | MAP domain-containing protein |
| WP_000868999.1 | -1.12 | 1.3E-04 | spoVG | septation regulator SpoVG |
| WP_000549278.1 | -1.12 | 9.4E-05 | essC | type VII secretion protein EssC |
| WP_001006445.1 | -1.11 | 4.1E-03 | norC | multidrug efflux MFS transporter NorC |
| WP_000111118.1 | -1.10 | 6.5E-03 |  | FeoB-associated Cys-rich membrane protein |
| WP_001165058.1 | -1.10 | 2.4E-02 |  | protein VraC |
| WP_000974460.1 | -1.10 | 7.9E-04 |  | organic hydroperoxide resistance protein |
| WP_000221958.1 | -1.09 | 7.1E-04 | leuB | 3-isopropylmalate dehydrogenase |
| WP_001802886.1 | -1.09 | 1.1E-03 |  | YagU family protein |
| WP_000249804.1 | -1.08 | 5.8E-03 | arcA | arginine deiminase |
| WP_001240826.1 | -1.08 | 8.2E-03 | esxA | WXG100 family type VII secretion effector EsxA |
| WP_000030812.1 | -1.07 | 4.1E-03 | purM | phosphoribosylformylglycinamide cyclo-ligase |
| WP_000247465.1 | -1.07 | 4.6E-03 |  | CHAP domain-containing protein |
| WP_001198024.1 | -1.07 | 4.3E-02 |  | TM2 domain-containing protein |
| WP_000077318.1 | -1.07 | 2.1E-03 |  | bifunctional homocysteine<br>S-methyltransferase/methylenetetrahydrofolate reductase |
| WP_000238664.1 | -1.07 | 6.6E-03 | purN | phosphoribosylglycinamide formyltransferase |
| WP_000239545.1 | -1.07 | 6.6E-04 | lukS-PV | Panton-Valentine bi-component leukocidin subunit S |
| WP_000686168.1 | -1.06 | 9.4E-05 |  | NAD(P)/FAD-dependent oxidoreductase |
| WP_001028431.1 | -1.06 | 2.3E-03 |  | NADPH-dependent FMN reductase |
| WP_000700921.1 | -1.06 | 4.9E-04 |  | carboxylesterase/lipase family protein |
| USA300HOU_RS00925 | -1.06 | 6.1E-03 |  | acyl-CoA/acyl-ACP dehydrogenase |
| WP_001092004.1 | -1.05 | 7.6E-04 |  | DUF2294 domain-containing protein |
| WP_000037332.1 | -1.05 | 5.4E-03 | cap8B | type 8 capsular polysaccharide synthesis protein Cap8B |
| WP_000648118.1 | -1.05 | 3.0E-04 |  | YtxH domain-containing protein |
| WP_000072278.1 | -1.04 | 7.9E-04 |  | sensor histidine kinase KdpD |
| WP_001045421.1 | -1.04 | 2.5E-02 |  | alpha/beta hydrolase |
| WP_000634175.1 | -1.04 | 8.5E-04 |  | universal stress protein |
| WP_000421410.1 | -1.03 | 5.0E-04 | xpt | xanthine phosphoribosyltransferase |
| WP_000709291.1 | -1.02 | 1.1E-02 | purH | bifunctional phosphoribosylaminoimidazolecarboxamide formyltransferase/IMP cyclohydrolase |
| WP_001146763.1 | -1.02 | 5.6E-03 |  | carboxymuconolactone decarboxylase family protein |
| WP_001101908.1 | -1.02 | 3.4E-03 | purD | phosphoribosylamine--glycine ligase |
| WP_001052483.1 | -1.02 | 1.2E-02 |  | hypothetical protein |
| WP_000691541.1 | -1.01 | 2.7E-02 |  | S8 family serine peptidase |
| WP_000840811.1 | -1.00 | 8.6E-03 |  | DUF1641 domain-containing protein |

**Table S3:** USA300 MRSA upregulated genes after exposure to *D. incerta*. Genes with adjusted p val < 0.05 and fold-change >2 are included. Rows are sorted by fold change.

| RefSeqID | Log<sub>2</sub>(FC) | adj p val | Gene Symbol | Protein |
| --- | --- | --- | --- | --- |
| WP_001790708.1 | 2.68 | 4.2E-02 |  | hypothetical protein |
| WP_000616842.1 | 2.57 | 1.8E-04 |  | ABC transporter ATP-binding protein |
| WP_000070866.1 | 2.08 | 2.6E-06 |  | Dps family protein |
| WP_001178619.1 | 2.06 | 9.8E-05 |  | solute carrier family 23 protein |
| WP_000066062.1 | 2.04 | 4.4E-04 | argH | argininosuccinate lyase |
| WP_001789981.1 | 2.04 | 4.6E-02 |  | hypothetical protein |
| WP_000754443.1 | 1.97 | 9.5E-05 |  | siderophore ABC transporter substrate-binding protein |
| WP_000660045.1 | 1.87 | 3.6E-03 |  | argininosuccinate synthase |
| WP_000932073.1 | 1.82 | 1.6E-02 |  | ferrous iron transport protein A |
| WP_000627551.1 | 1.75 | 1.2E-02 |  | DUF3969 family protein |
| WP_000003870.1 | 1.67 | 3.3E-05 | pyrR | bifunctional pyr operon transcriptional regulator/uracil phosphoribosyltransferase PyrR |
| WP_001016166.1 | 1.63 | 8.8E-05 |  | aspartate carbamoyltransferase catalytic subunit |
| WP_001245577.1 | 1.62 | 9.7E-03 |  | iron chelate uptake ABC transporter family permease subunit |
| WP_000213873.1 | 1.61 | 1.8E-05 |  | DHA2 family efflux MFS transporter permease subunit |
| WP_000539688.1 | 1.59 | 3.1E-02 |  | DUF2951 domain-containing protein |
| WP_000793605.1 | 1.57 | 3.1E-04 | argF | ornithine carbamoyltransferase |
| WP_000738247.1 | 1.57 | 2.0E-04 |  | HlyD family efflux transporter periplasmic adaptor subunit |
| WP_000083807.1 | 1.56 | 3.7E-05 |  | PTS mannitol transporter subunit IICB |
| WP_000590809.1 | 1.54 | 1.1E-02 |  | amino acid ABC transporter ATP-binding protein |
| WP_000402904.1 | 1.53 | 3.5E-02 |  | hypothetical protein |
| WP_001040261.1 | 1.52 | 9.9E-03 |  | hypothetical protein |
| WP_000052781.1 | 1.52 | 3.1E-05 | ahpC | alkyl hydroperoxide reductase subunit C |
| WP_001074342.1 | 1.50 | 2.4E-04 | arcC | carbamate kinase |
| WP_000612128.1 | 1.49 | 2.5E-02 |  | urease subunit beta |
| WP_000432077.1 | 1.49 | 2.1E-04 |  | YfcC family protein |
| WP_000930486.1 | 1.49 | 7.4E-05 | ahpF | alkyl hydroperoxide reductase subunit F |
| WP_000649907.1 | 1.41 | 8.0E-03 |  | ABC transporter permease subunit |
| WP_000767028.1 | 1.40 | 3.0E-04 |  | dihydroorotase |
| USA300HOU_RS09550 | 1.40 | 9.2E-03 |  | hypothetical protein |
| WP_000876318.1 | 1.36 | 1.3E-02 |  | ABC transporter permease |
| WP_000828726.1 | 1.26 | 2.3E-03 |  | sodium-dependent transporter |
| WP_000549734.1 | 1.24 | 1.7E-02 | cidA | holin-like murein hydrolase modulator CidA |
| WP_000798979.1 | 1.23 | 8.7E-04 |  | PepSY domain-containing protein |
| WP_001549577.1 | 1.21 | 3.4E-03 |  | YjiH family protein |
| WP_000571584.1 | 1.18 | 8.5E-04 | gltS | sodium/glutamate symporter |
| WP_000064778.1 | 1.15 | 1.2E-04 |  | amino acid permease |
| WP_001200541.1 | 1.11 | 1.6E-02 | crcB | fluoride efflux transporter CrcB |
| WP_000181819.1 | 1.11 | 3.0E-02 |  | dUTP pyrophosphatase |
| WP_000210828.1 | 1.11 | 1.0E-03 | tdcB | bifunctional threonine ammonia-lyase/L-serine ammonia-lyase TdcB |
| WP_000755150.1 | 1.10 | 8.4E-03 |  | head-tail adaptor protein |
| WP_000985235.1 | 1.10 | 1.3E-03 |  | phage/plasmid primase, P4 family |
| WP_000447733.1 | 1.09 | 1.1E-04 | xerD | site-specific tyrosine recombinase XerD |
| WP_011447039.1 | 1.09 | 8.9E-04 |  | putative holin-like toxin |
| WP_000725225.1 | 1.08 | 1.2E-04 |  | CHAP domain-containing protein |
| WP_000414205.1 | 1.08 | 4.8E-03 |  | hypothetical protein |
| WP_001008722.1 | 1.08 | 3.8E-04 | uhpT | hexose-6-phosphate:phosphate antiporter |
| WP_001120199.1 | 1.06 | 3.5E-02 |  | DUF771 domain-containing protein |
| WP_001795631.1 | 1.05 | 2.5E-04 | tatC | twin-arginine translocase subunit TatC |
| WP_000789821.1 | 1.05 | 4.4E-02 | isdC | heme uptake protein IsdC |
| WP_000959424.1 | 1.05 | 2.2E-03 | ald | alanine dehydrogenase |
| WP_000861038.1 | 1.05 | 7.4E-03 |  | CHAP domain-containing protein |
| WP_000414685.1 | 1.04 | 1.3E-04 | norB | multidrug efflux MFS transporter NorB |
| WP_001549178.1 | 1.04 | 4.5E-02 |  | DUF2482 family protein |
| WP_001261987.1 | 1.00 | 1.9E-03 | rimM | ribosome maturation factor RimM |

**Table S4:**
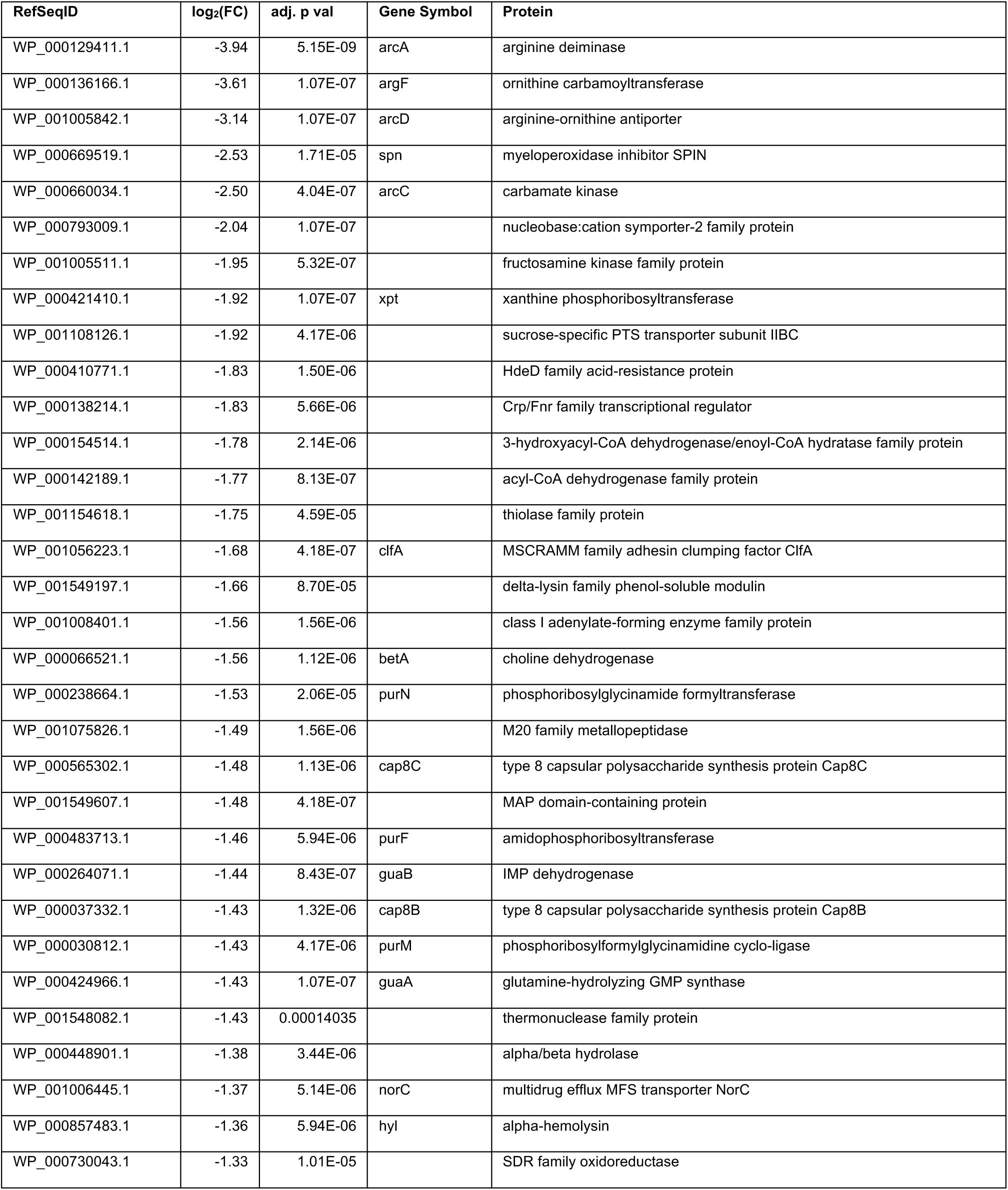
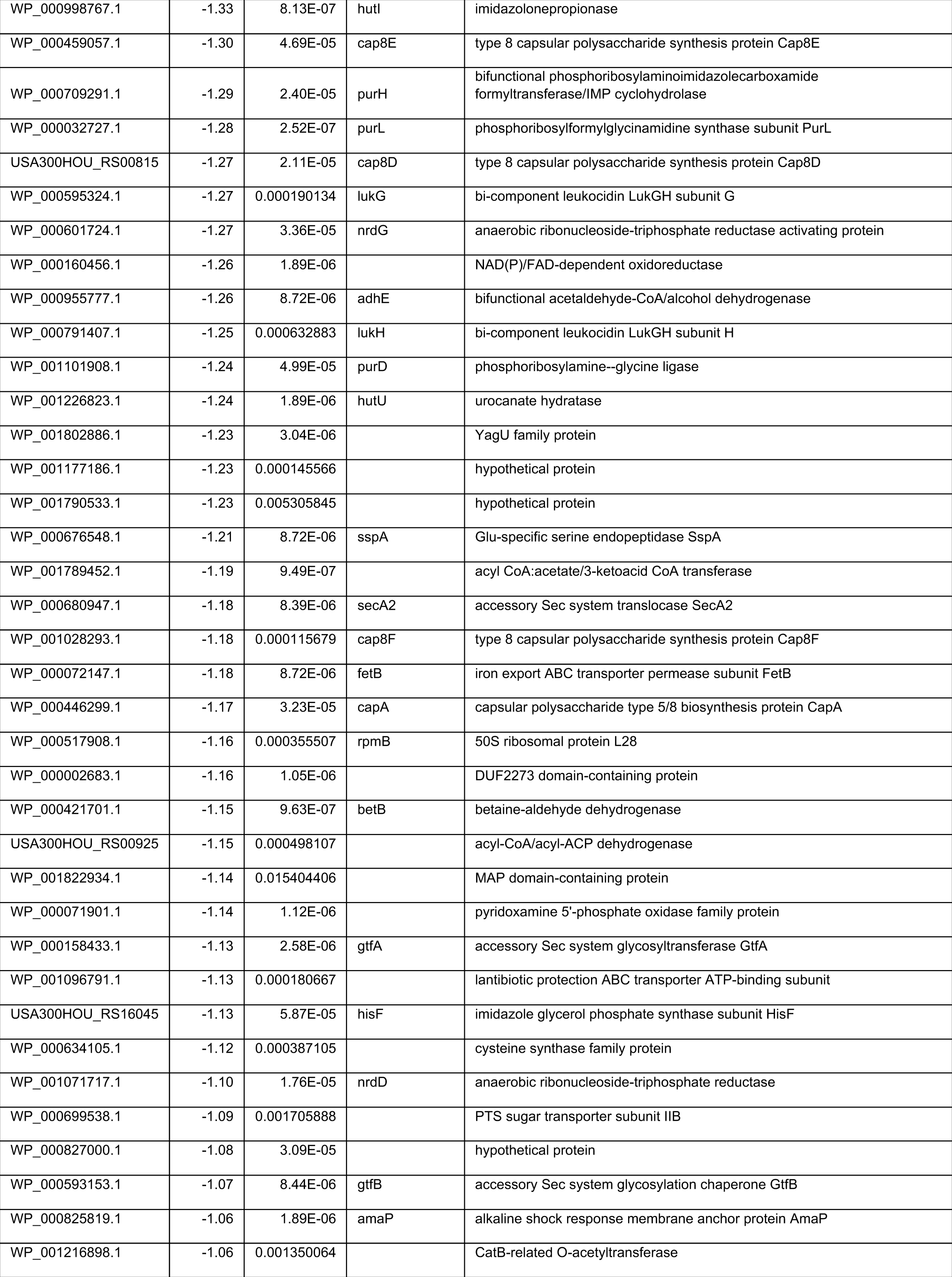
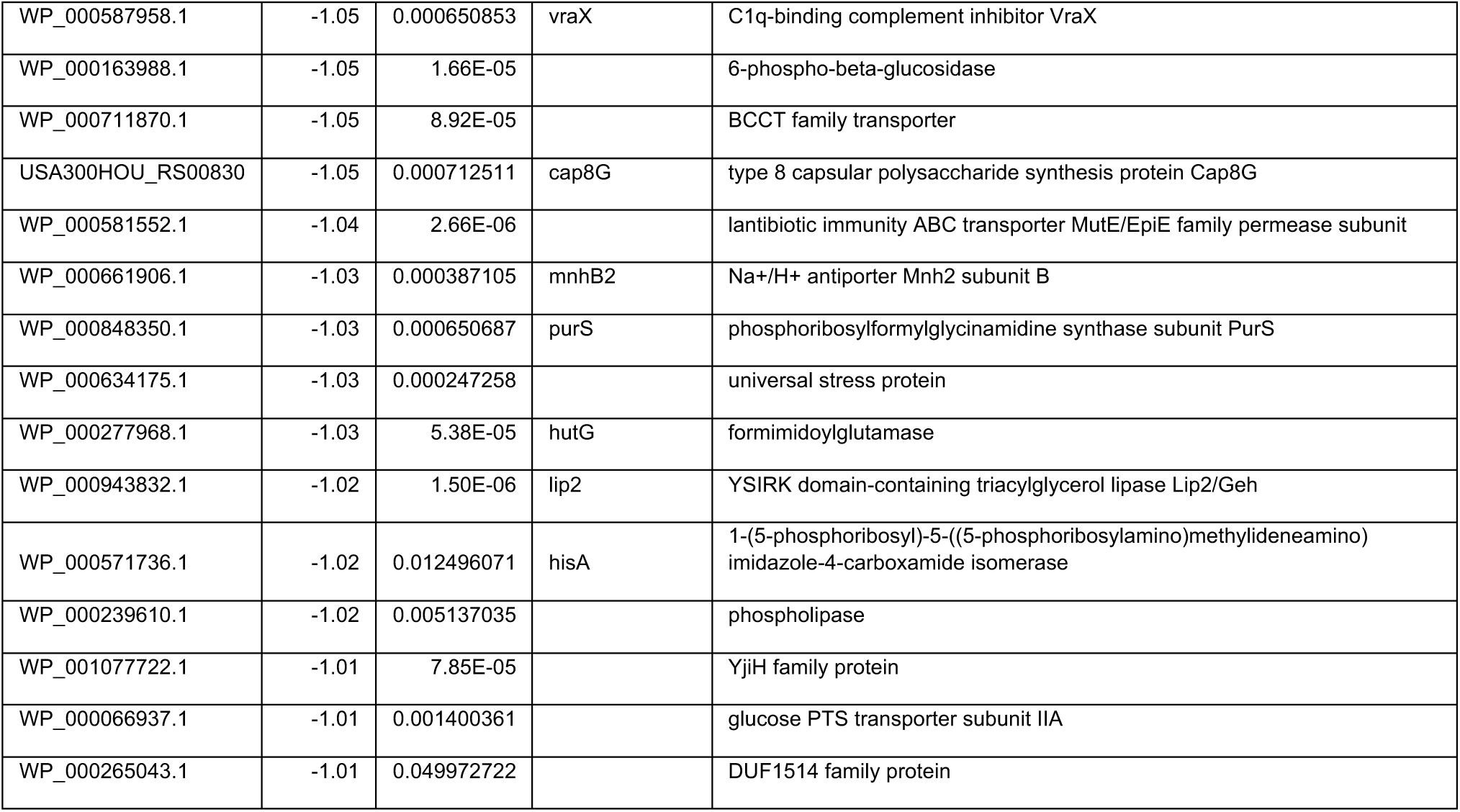
*S. aureus SA113* downregulated genes after exposure to *D. incerta*. Genes with adjusted p val < 0.05 and fold-change >2 are included. Rows are sorted by fold change.

**Table S5:**
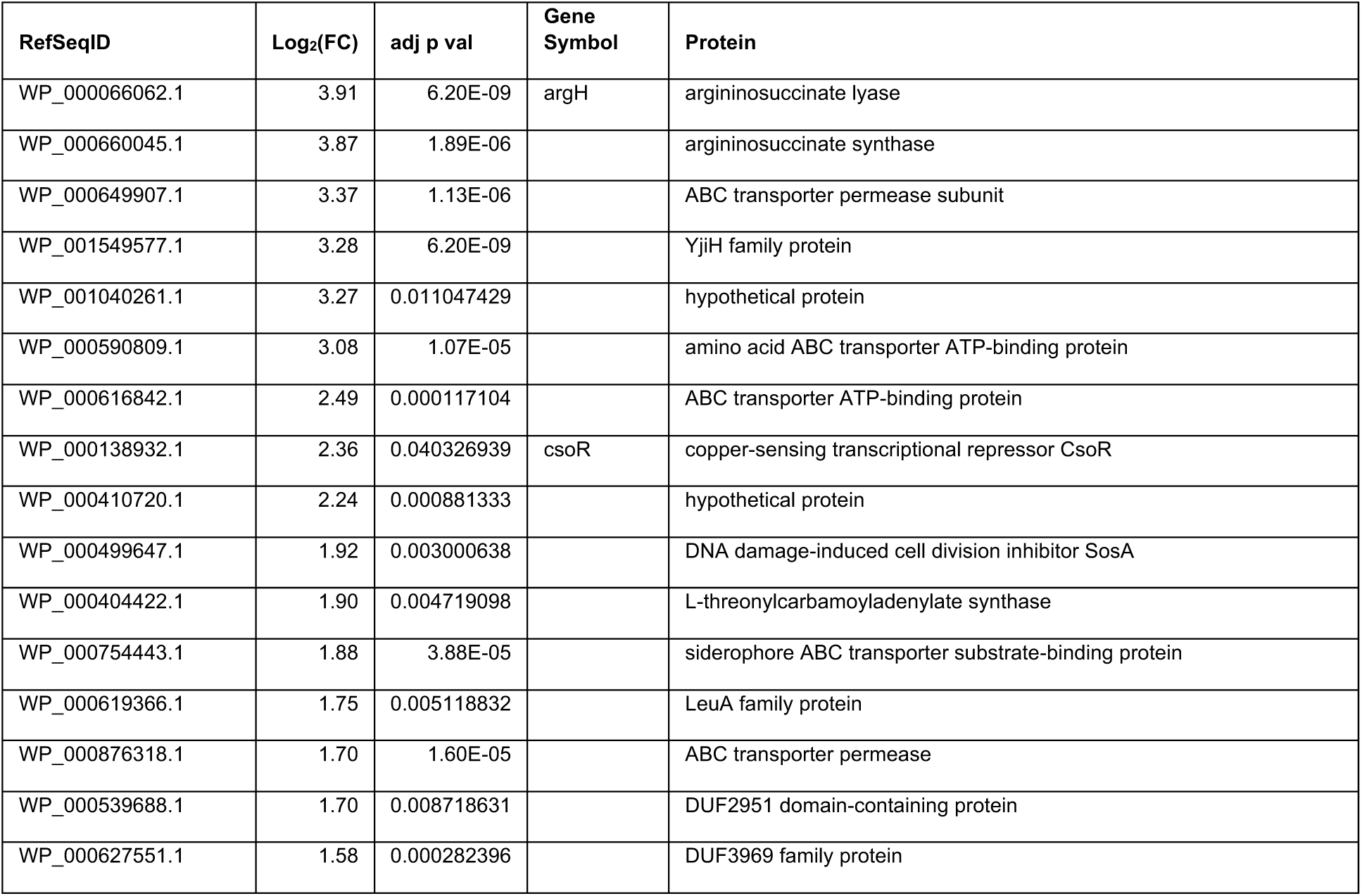
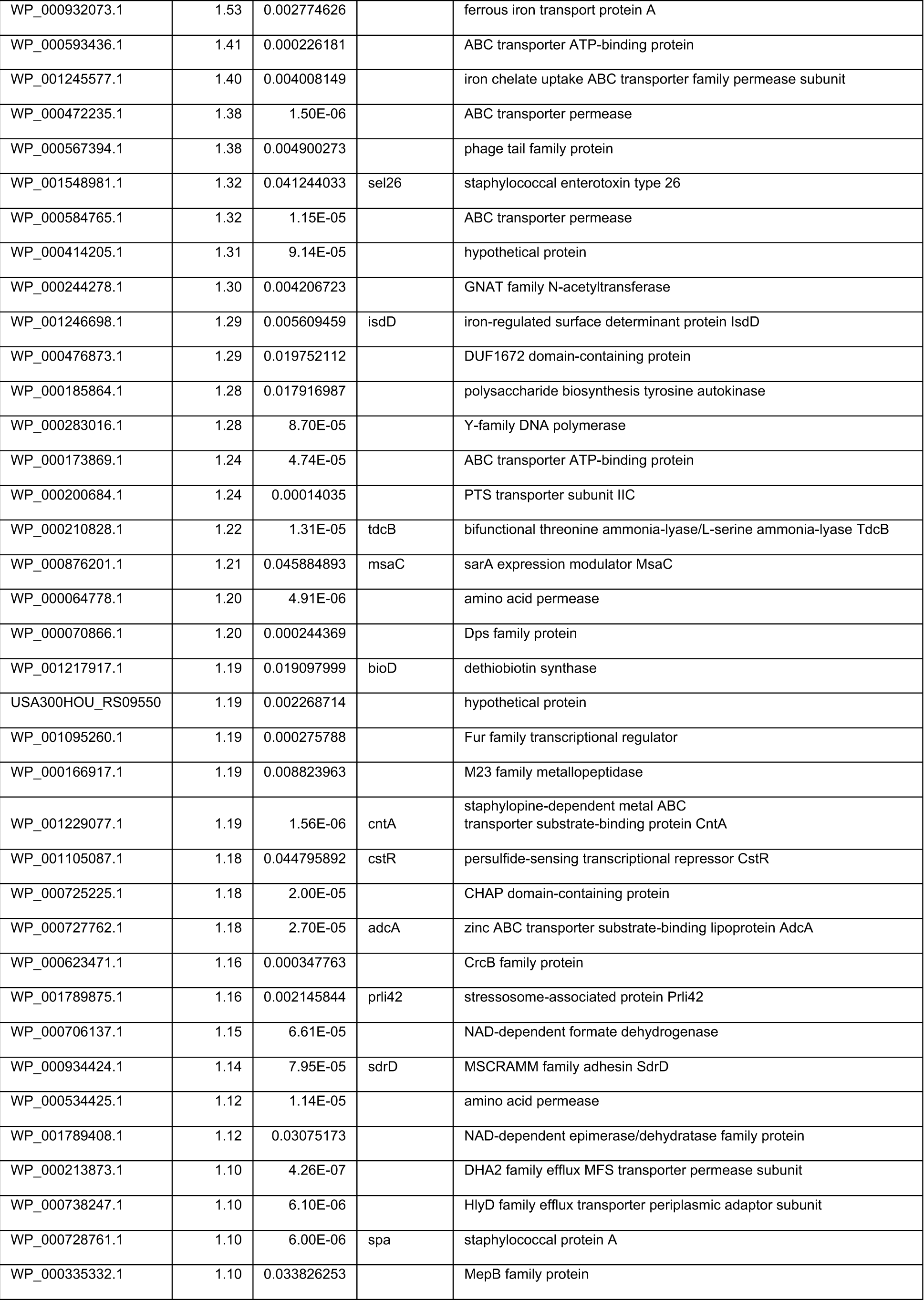
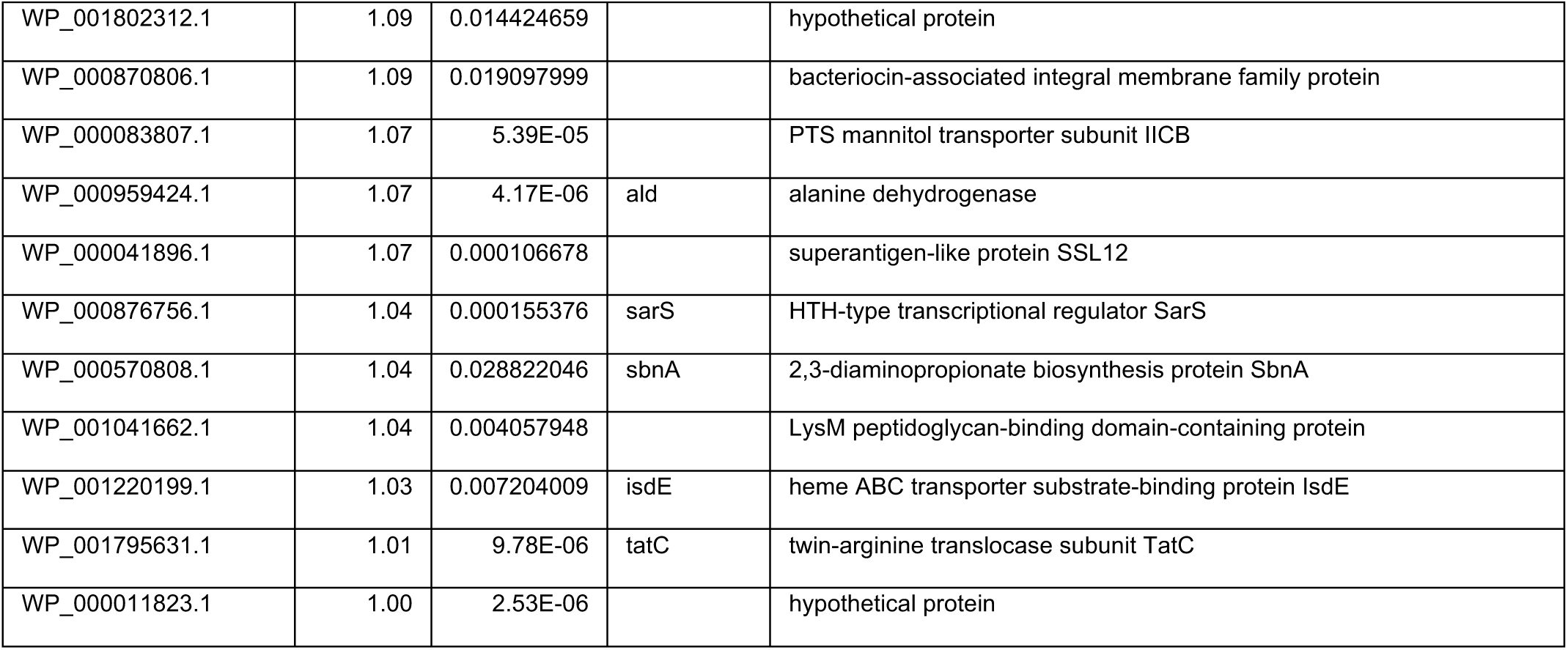
*S. aureus SA113* upregulated genes after exposure to *D. incerta*. Genes with adjusted p val < 0.05 and fold-change >2 are included. Rows are sorted by fold change.

**Figure S2:**
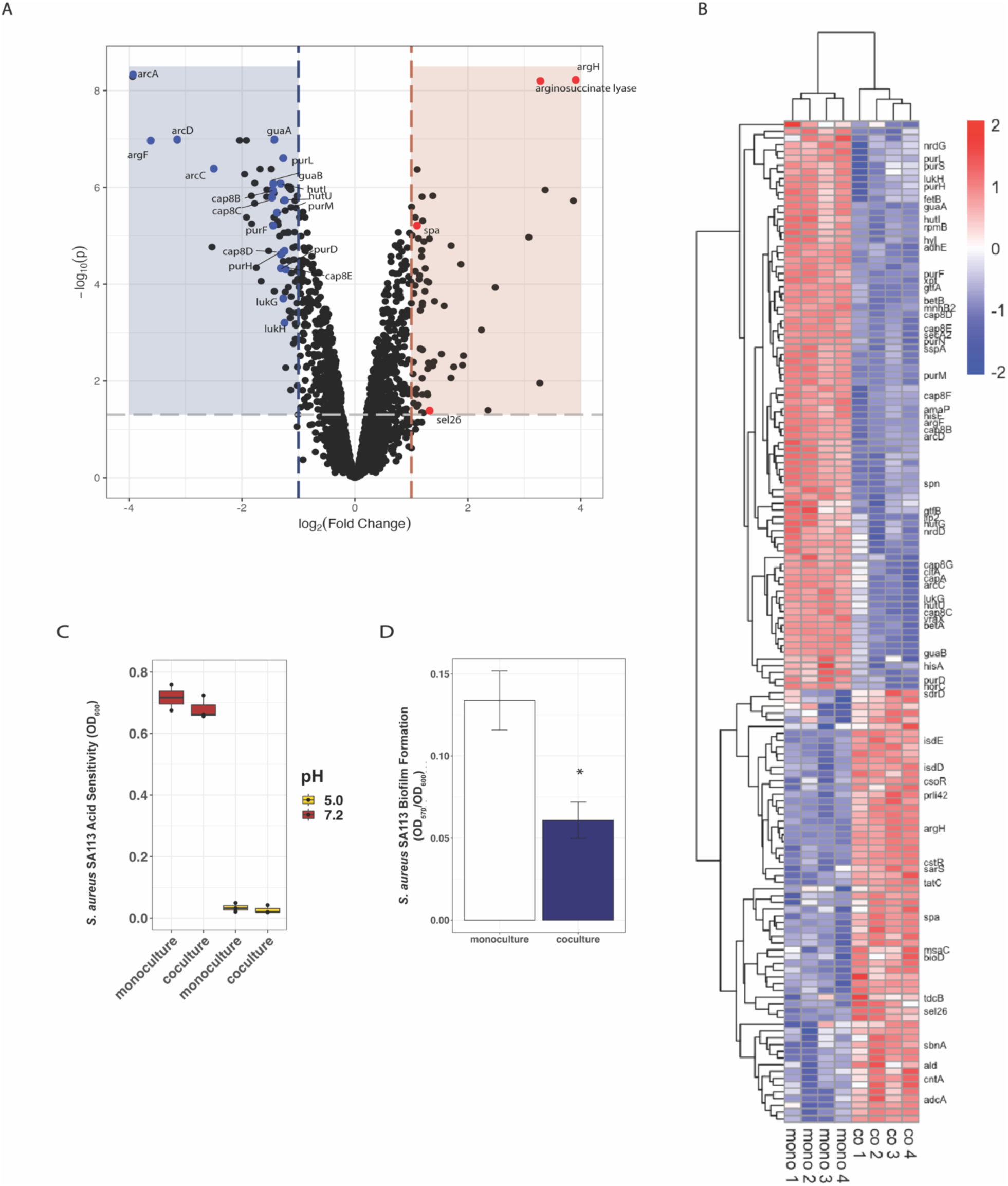
Transcriptional profiling of *S. aureus* SA113 reveals similar changes in gene expression and biofilm formation compared to USA300 MRSA. **A)** Volcano plot of genes in *S. aureus* SA113 that were differentially expressed during *D. incerta* coculture compared to monoculture. Shaded areas highlight genes whose adjusted p < 0.05 and fold change > 2.0 (blue – downregulated, red-upregulated). Bold points mark genes of interest, which were found within operons containing multiple differentially expressed genes. **B)** Heatmap depicting differentially expressed genes in USA300 MRSA with adjusted p< 0.05 and fold change >2.0. For visualization purposes, genes were ordered by hierarchical clustering based on Pearson correlation. Values were scaled to the row mean. **C)** *S. aureus* SA 113 cell density in response to acidified media (pH = 5.0, yellow) compared to neutral pH media (red) and coculture with *D. incerta*. 3 wells per condition were used. **D)** *S. aureus* SA113 biofilm formation measured via crystal violet stain retention (OD_570_) normalized to cell growth (OD_600_). 4 wells per condition were used. T-test, * p < 0.05

